# Progressive recruitment of distal MEC-4 channels determines touch response strength in *C. elegans*

**DOI:** 10.1101/587014

**Authors:** S. Katta, A. Sanzeni, A. Das, M. Vergassola, M.B. Goodman

## Abstract

Touch deforms, or strains, the skin beyond the immediate point of contact. The spatiotemporal nature of the touch-induced strain fields depend on the mechanical properties of the skin and the tissues below. Somatosensory neurons that sense touch branch out within the skin and rely on a set of mechano-electrical transduction channels distributed within their dendrites to detect mechanical stimuli. Here, we sought to understand how tissue mechanics shape touch-induced mechanical strain across the skin over time and how individual channels located in different regions of the strain field contribute to the overall touch response. We leveraged *C. elegans’* touch receptor neurons (TRNs) as a simple model amenable to *in vivo* whole-cell patch clamp recording and an integrated experimental-computational approach to dissect the mechanisms underlying the spatial and temporal dynamics that we observed. Consistent with the idea that strain is produced at a distance, we show that delivering strong stimuli outside the anatomical extent of the neuron is sufficient to evoke MRCs. The amplitude and kinetics of the MRCs depended on both stimulus displacement and speed. Finally, we found that the main factor responsible for touch sensitivity is the recruitment of progressively more distant channels by stronger stimuli, rather than modulation of channel open probability. This principle may generalize to somatosensory neurons with more complex morphologies.

**Summary:** Through experiment and simulation, Katta *et al*. reveal that pushing faster and deeper recruits more and more distant mechano-electrical transduction channels during touch. The net result is a dynamic receptive field whose size and shape depends on tissue mechanics, stimulus parameters, and channel distribution within sensory neurons.

## Introduction

Sensory receptor neurons are classified by their receptive fields -- the region of their sensory space in which a stimulus elicits a response, such as a spot in the visual field or a frequency of sound. The receptive field of a somatosensory neuron is a combined function of its dendritic arbor and the mechanical strain, or deformation, induced by touch. Mechanical strain decreases with distance from the stimulus; the extent of the strain field and the speed of propagation depend on the intensity of the stimulus and the material properties of the neurons and skin being touched. With light and chemical stimuli, a single photon or ligand can only activate a single receptor protein at a time, even if many are in close proximity. With touch, a single indentation will simultaneously affect all eligible receptors that fall within the strain field, to a degree dependent on their distance from the point of stimulation. In the case of somatosensation, this means that multiple channels at spatially distinct sites within a neuron, or even multiple neurons, are liable to be activated to varying degrees by the same stimulus.

Intricate anatomical structures dedicated to mechanosensation are present in worms, flies, mice, cats, and humans. From the chordotonal organs and campaniform receptors of insects to Merkel cell touch domes and low-threshold mechanoreceptors arrayed around hair follicles in mammals, these structures place mechanosensitive proteins in specific arrangements around features of the skin or other mechanosensory organs (Katta et al., 2015). Increasing anatomical and functional data have provided insights into how these structures work. In many cases the complexity of skin has made it difficult to understand the strain field. While useful models have been created (Lesniak and Gerling, 2009; Quindlen et al., 2015; Quindlen-Hotek and Barocas, 2018; Sanzeni et al., 2018), few studies build on these models to generate an integrated understanding of how mechanical strain is detected by mechanosensitive proteins, including ion channels, distributed within somatosensory neurons.

Here, we probed how mechano-electrical transduction (MeT) channel distribution interacts with the strain field of a mechanical stimulus in a *C. elegans* touch receptor neuron (TRN). This system is a useful model because the factors governing the strain field are straightforward and the arrangement of channels is simple and well-characterized. Furthermore, we can directly measure the mechanoreceptor currents (MRCs). The six *C. elegans* TRNs (ALMR/L, PLMR/L, AVM, and PVM) extend long, unusually straight neurites (Krieg et al., 2017) that are embedded in the epidermal cells that form the worm’s skin or cuticle (Chalfie and Sulston, 1981; Chalfie and Thomson, 1979). External mechanical loads evoke MRCs that are carried by MEC-4-dependent MeT channels and are activated at both the application and removal of mechanical stimuli (O’Hagan et al., 2005). The MEC-4 protein belongs to a large superfamily of non-voltage-gated ion channels conserved in animals but absent from microbes and plants (Katta et al., 2015). MEC-4 proteins localize to discrete puncta that are arrayed along the entire length of TRN neurites (Chelur et al., 2002; Cueva et al., 2007; Emtage et al., 2004). These anatomical properties make it easier to interpret how the geometry of the strain field depends on the size and speed of a given stimulus and determine which subset of MeT channels is likely to be affected by each stimulus.

Because MRC amplitude depends more on indentation than it does on applied force (Eastwood et al., 2015), we developed and deployed a stimulator system enabling fast indentation and concurrent optical monitoring of probe movement. We used this system to measure MRCs evoked by mechanical stimuli delivered inside and outside of the anatomical receptive field of the ALM neuron and as a function of indentation depth and speed. By combining these experimental results with simulations (Sanzeni et al., 2018), we identified a biophysical mechanism linking indentation to MRCs. Specifically, we show that MRC size and time-course depend on both the modulation of open probability and the recruitment of distal channels in a model somatosensory neuron.

## Materials and Methods

### Nematode Strains

We used three strains of transgenic *C. elegans* nematodes: TU2769 *uIs31*[*mec-17p::gfp*] *III* (O’Hagan et al., 2005) for electrophysiology, GN865 *uIs31*[*mec-17p::GFP*] *III; kaIs12*[*col-19::GFP*] for imaging cuticle indentation, and GN753 *pgSi116*[*mec-17p::mNeonGreen::3xFLAG::MEC-4::tbb-2 3’UTR*] *II* for visualizing MEC-4 puncta. The *uIs31* transgene is a TRN-specific GFP marker that enables us to conduct *in vivo* recordings in TRNs. The *kaIs12* transgene encodes a GFP fusion to a collagen that labels cuticular annuli (Thein et al., 2003). The *pgSi116* transgene expresses MEC-4 tagged with mNeonGreen in the touch receptor neurons to visualize MEC-4 puncta. We grew animals on OP50 and used well-fed subjects as late-L4 larvae or young adults. On average, the animals used for electrophysiology had a body length of 1298 ± 93 µm (mean ± SD, *N* = 53 with a range of 1064 to 1547) long and diameter of 39 ± 4 µm (with a range of 32 to 50) at the terminal bulb of the pharynx.

### Imaging Cuticle Deformation

We immobilized GN865 worms using 2% agarose pads with WormGlu (GluStitch); subjects were either left intact or dissected as described previously (Eastwood et al., 2015; O’Hagan et al., 2005). We used an Orca Flash 4.0LTv2 camera (Hamamatsu) controlled by μManager (Edelstein et al., 2010) on an upright microscope (E600FN, Nikon) under a 60X objective for imaging.

### Electrophysiology

We recorded from the ALMR and ALML touch receptor neurons in TU2769 worms. Due to the geometric constraints of this stimulator system, we recorded from ALMR when stimulating anterior to the cell body and from ALML when stimulating posterior to the cell body. These two neurons are bilaterally symmetric and exhibit no detectable variation in voltage- or touch-evoked currents (not shown). The extracellular solution contained (in mM): NaCl (145), KCl (5), MgCl_2_ (5), CaCl_2_ (1), and Na-HEPES (10), adjusted to a pH of 7.2 with NaOH. Before using the solution, we added 20mM of D-glucose, which brought the osmolarity to ∼325mOsm. The intracellular solution contained (in mM): K-gluconate (125), KCl (18), NaCl (4), MgCl_2_ (1), CaCl_2_ (0.6), K-HEPES (10), and K_2_EGTA (10), adjusted to a pH of 7.2 with KOH. Before use, we added 1mM of sulforhodamine 101 (Invitrogen) to help visualize whether the neuron was being successfully recorded.

We used an EPC-10 USB amplifier controlled by Patchmaster software (version 2×90.1, HEKA/Harvard Biosciences) to set the membrane voltage and control the mechanical stimulator. We corrected the membrane voltage for the liquid-junction potential (−14 mV) between the extracellular and intracellular solutions; we also corrected for errors that may have resulted from an imperfect space clamp based on stimulus distance, as described previously (Goodman et al., 1998; O’Hagan et al., 2005). We filtered the analog data at 2.9 kHz and digitized it at 10 kHz.

### Mechanical Stimulation

Some of the previous electrophysiological studies on *C. elegans* mechanoreceptor neurons have used open-loop systems with a piezoelectric bimorph (Árnadóttir et al., 2011; Bounoutas et al., 2009; Chen and Chalfie, 2015; Geffeney et al., 2011; O’Hagan et al., 2005) or a piezoelectric stack with no independent measurement of stimulator motion (Kang et al., 2010). We previously employed a slow closed-loop system driven by a piezoelectric stack with a piezoresistive cantilever for force detection (Eastwood et al., 2015). Here, we used an open-loop system adapted from the piezoelectric stack system with a photodiode motion detector described by Peng and Ricci (2016) in their study on hair cells. This system enables faster stimulation than the force-clamp system and allows us to measure the time course of stimulation.

Using marine epoxy (Loctite), we attached an open-loop, piezoelectric stack actuator (PAS-005, ThorLabs, 20 µm travel) to a tungsten rod (L = 8 in.) to damp vibration. We mounted the rod-mounted actuator on a micromanipulator for positioning (MP-225, Sutter) and controlled the piezoelectric stack via an analog voltage output on a HEKA EPC-10 PLUS amplifier and Patchmaster software. We filtered this analog signal at 2.5 kHz for steps or 5 kHz for ramps and sines on an eight-pole Bessel filter (LPF-8, Warner Instruments) and used a high-voltage, high-current amplifier (Peng and Ricci, 2016) to achieve a signal between 0-75 V. The stack was biased with a starting offset of 3-4 μm, and the largest displacement we used was 3-4 μm short of the 20 μm total travel limit. This protocol ensured that stack motion was linearly related to the analog voltage signal.

Beads glued to force-clamp cantilevers create a defined and reproducible contact surface that is easy to model (Eastwood et al., 2015; Petzold et al., 2013). We adapted this technique for a system relying on stiff glass, rather than silicon probes and for axial stimulus motion. We pulled and polished borosilicate glass pipettes (Sutter, BF150-86-10) to a tip diameter that was half or three-fourths of the diameter of the bead that we intended to use. We attached black polyethylene beads (20-24 μm diameter; BKPMS-1.2, Cospheric) to the glass pipettes using UV-curable glue (Loctite 352, Henkel). The bead-attached pipettes were then waxed in place to a 3D-printed acrylic pipette holder (custom design available from https://3dprint.nih.gov as Model ID 3DPX-010770) and epoxied to a steel tip (PAA001, ThorLabs) that was mounted on the piezo stack.

Step protocols had a 250 ms or 300 ms hold at the commanded indentation, and used an inter-stimulus interval (ISI) of 1 s. Trapezoidal protocols had a 300 ms hold and 2 s ISI. Sinusoidal stimulus profiles consisted of a 1-s sinusoidal stimulus epoch flanked by 300 ms steps, and 2 s ISI.

### Motion Detection

To monitor the motion of the stimulus probe, we modified the system reported by Peng and Ricci (2016) to detect larger movements by using the SPOT-2D segmented photodiode (OSI Optoelectronics) and a higher resistance in the differential amplifier circuit. The photodiode was mounted in an XY translator on top of a rotation stage (ST1XY-D, LCP02R, ThorLabs) such that the photodiode gap was aligned perpendicular to the direction of the probe motion. This apparatus was fixed above a secondary camera port on the microscope (Eclipse E600FN, Nikon) with no additional magnification.

Before obtaining an on-cell, high resistance seal for patch clamp, we aligned the front edge of the bead under the 60X objective with the highest clearly visible edge of the worm’s cuticle. After achieving whole-cell access, we moved the microscope to visually align the image of the bead with the outer edge of the photodiode. As the bead moved, the output from the photodiode circuit was read directly by Patchmaster, and we used this signal to check that resonance during steps and relative attenuation of sine amplitude at high frequencies were small.

At the end of each recording, we calibrated the image motion to actual motion. We drove the XY translator in steps of known distances in a direction that was opposite to the probe motion using a stepper motor actuator (ZFS-06, ThorLabs) controlled by Kinesis software (ThorLabs). We took these steps once with the probe at the edge of the worm and once with only the worm. Distances were measured relative to this starting position. The worm-only trace was subtracted from the probe+worm trace to obtain the relationship between the motion of the probe image and the known movement of the photodiode. We corrected stimulus traces from the recording using a piecewise cubic polynominal interpolation method (’pchip’, MATLAB R2015b) of the mean voltage during calibration steps vs. the commanded photodiode motion at each step.

### Data Analysis

We measured whole-cell capacitance and series resistance as previously described (Goodman et al., 1998). We conducted data analysis using MATLAB from Mathworks (data import and analysis functions are available online at: https://github.com/wormsenseLab/Matlab-PatchMaster/tree/vDynamicsJGP) and Igor Pro 6.37 (Wavemetrics).

Our analysis only included recordings with a holding current less than −10 pA at −60 mV and a series resistance less than 250 MΩ. As these experiments were conducted at constant voltage, the voltage errors due to uncompensated series resistance were negligible, and we did not correct the reported data for these errors. Where noted, we corrected for voltage attenuation due to space-clamp errors as previously described (O’Hagan et al., 2005). Briefly, we computed the theoretical capacitance for each neuron based on body size and compared it to the measured capacitance to estimate the ratio of membrane resistance to axial resistance for ALM. This allowed us to calculate the length constant for each neuron and estimate the actual voltage at the center of the stimulus, which we used as the average channel location for the purpose of accounting for space-clamp error. We calculated total charge, or the area under the MRC curve, from stimulus start to 150-ms after stimulus end, and subtracted the charge at baseline for an equivalent time period.

### MEC-4 puncta imaging and data analysis

We mounted young adult GN753 *pgSi116 mNeonGreen::3xFLAG::MEC-4* transgenic animals on grooved 5% agarose pads dissolved in M9 buffer. Grooved pads were prepared by casting agarose on the surface of an LP vinyl record (Rivera Gomez and Schvarzstein, 2018). Animals were immobilized using levamisole (5 mM) dissolved in M9 buffer. Micrographs were acquired with an automated inverted epifluorescence microscope (Keyence BZ-X800) equipped with software and hardware to generate stitched images of entire ALM neurons with a 40x objective (Nikon plan apo 40x/1.0 oil).

The stitched images were pre-processed using ImageJ, where we manually traced a line along the neurite from the cell body to the distal tip to create a 20-pixel wide straightened image of the neurite. We then used these straightened neurite images to analyze MEC-4 puncta distribution using a custom Python script (https://github.com/wormsenseLab/Puncta_analysis/tree/vJGP).

Briefly, we averaged the pixel intensities in the middle rows to calculate the raw neurite fluorescence, subtracted the average background fluorescence at the top and bottom, divided by the background to normalize across animals, and smoothed with a 2-pixel maximum filter to reduce noise. Peaks in the fluorescence traces were detected automatically (‘find_peaks’, scipy library). Inter-punctum intervals were calculated by measuring the distance between adjacent peaks along the neurite and the location of each inter-punctum interval is calculated as the distance of the mid-point between the adjacent peaks from the cell body.

### Simulations

For simulations, we used the model described in Sanzeni *et al*. (2018) and model parameters derived by fitting the dataset from Eastwood *et al*. and the FALCON closed-loop stimulation system (Eastwood et al., 2015 and Table 1). We took this approach for the following reasons. First, the previous dataset directly linked force, indentation, and current, enabling the inference of model parameters. By contrast, the current data set was collected using a mechanical stimulation system that enables high-speed indentation at the expense of this direct link. Second, single channel parameters are, to a first approximation, preserved across worms and independent of internal pressure (Sanzeni et al., 2018). Consequently, we use single channel parameters derived from the dataset of Eastwood *et al*. (2015), which allows us to infer internal pressure, and show model predictions obtained using either a typical value of 4kPa internal pressure (Sanzeni et al., 2018) or different values of internal pressure (e.g. Fig. 3). The stimulator was modeled as a rigid sphere with a 20 μm diameter, in accord with the stimulation apparatus used in this study.

**Table 1.** Modeling parameters for simulation. Parameters were derived from fitting the model to the dataset from Eastwood *et al*. (2015), as described in Sanzeni *et al*. (2018). We show parameter sets derived from fitting based on the 1.4μm inter-punctum interval (IPI) used in the majority of the paper, as well as with longer IPIs. *τ* represents the time constant of relaxation for the linker, while *g*_0_ and *g*_1_ are dimensionless parameters relating linker elongation to the free energy of the channel. The reaction rate for transitioning from C ⇋ S is controlled by a base rate *r*_*c*P*s*_ and a factor *b* that controls how dependent the final rate is on the free energy change (Sanzeni et al., 2018, Appendix L). Similarly, *r*_*s*P*o*_ and *d* control the reaction rate for the S ⇋ O transition. *α* is the ratio of the free energy difference between C and S over the free energy difference between C and O. By combining these parameters with the recording conditions, we can calculate the ratio of the single-channel current in the subconductance state (*i*_*s*_) over the current carried by a fully open channel (*i*_*o*_). The strain energy density was not a fitted parameter, but was derived from single channel dynamics as discussed in the Materials and Methods section, and represents the energy available to the channel as a result of deformation due to step stimulation.

| Inter-punctum interval ($\mu\text{m}$) | 1.4 | 2 | 2.5 | 3 | 3.5 | 4 |
| --- | --- | --- | --- | --- | --- | --- |
| $\tau$ (ms) | 2.96 | 2.62 | 2.33 | 2.28 | 1.99 | 2.00 |
| $g_0$ | 9.17 | 8.85 | 8.77 | 8.70 | 9.35 | 9.71 |
| $g_1$ | -3058 | -2950 | -4386 | -4348 | -4673 | -4854 |
| $1/r_{cs}$ (ms) | 22.0 | 22.4 | 31.7 | 17.6 | 35.7 | 39.7 |
| $b$ | -0.869 | -0.852 | -0.742 | -0.814 | -0.694 | -0.677 |
| $1/r_{so}$ (ms) | 470 | 350 | 317 | 258 | 397 | 463 |
| $d$ | -0.194 | -0.247 | -0.262 | -0.332 | -0.304 | -0.31 |
| $\alpha$ | 0.458 | 0.466 | 0.459 | 0.482 | 0.511 | 0.52 |
| $i_s/i_o$ | 0.216 | 0.287 | 0.348 | 0.32 | 0.469 | 0.527 |
| Strain energy density (kPa) | 7.69 | 4.22 | 2.99 | 2.19 | 1.34 | 0.87 |

As in our recent computational study (Sanzeni et al., 2018), we built a finite element mechanical model of the worm as a pressurized, thin-shelled cylinder. A three-state (closed, sub-conducting, open) gating model of the channels allowed us to estimate the probabilities for individual channels to open. We opted to include the sub-conducting state because single channel recording of heterologously expressed MEC-4-dependent channels demonstrate a subconducting state (Brown et al., 2007). Additionally, including the sub-conducting state improved the match between simulated and experimental data (Eastwood et al., 2015). The total current flowing along the TRN is the sum of the currents flowing through the channels arrayed along the neuron that spans the length of the worm. In simulations, we used a fixed inter-punctum interval of 1.4µm and assumed that each punctum contained a single functional MeT channel. These choices are supported by empirical studies of endogenous channel distribution and estimates of the number of functional channels per punctum (Cueva et al., 2007).

The Strain Energy Density (SED) needed to activate a single channel was estimated as follows. We considered a single channel located 1μm below the cuticle surface and 1μm from the stimulus center and computed its current response to step stimuli of different displacements. For any given displacement *w*, we also computed the SED produced at the channel position *x* (Audoly and Pomeau, 2010) as

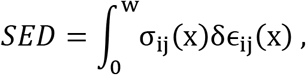

where *σ*_*ij*_ and *δ*∊_*ij*_ are the stress tensor and the change in strain tensor, respectively. Using this approach and assuming that reliable neuronal activation would be achieved following stimuli sufficient to achieve a half-maximal MRC amplitude, we estimated the SED needed to activate a touch receptor neuron.

## Results

Mechanoreceptor currents in *C. elegans* touch receptor neurons (TRNs) are carried by MeT channels distributed in puncta along the length of each sensory neurite (Chelur et al., 2002; Cueva et al., 2007; Emtage et al., 2004). For many other types of channels, an increase in macroscopic current is thought to reflect an increase in the open probability of a population of channels that receive the same stimulus at the same time. For MeT channels, however, an increase in the macroscopic current cannot be interpreted solely as an increase in open probability. As local indentation deforms tissue at distal sites (Fig. 1A-1C) and in a manner that depends on indentation depth (Elmi et al., 2017; Sanzeni et al., 2018), stimulus intensity governs how many channels are reachable by the stimulus and the probability that a given channel will open (Fig. 1A). Similarly, we expect that stimulus speed influences the evolution of the strain field, thereby introducing an additional factor that affects the synchrony or phase of MeT channel activation. As such, the spatial distribution of these MeT channels can affect how they respond to stimuli that vary across space and time.

**Figure 1.**
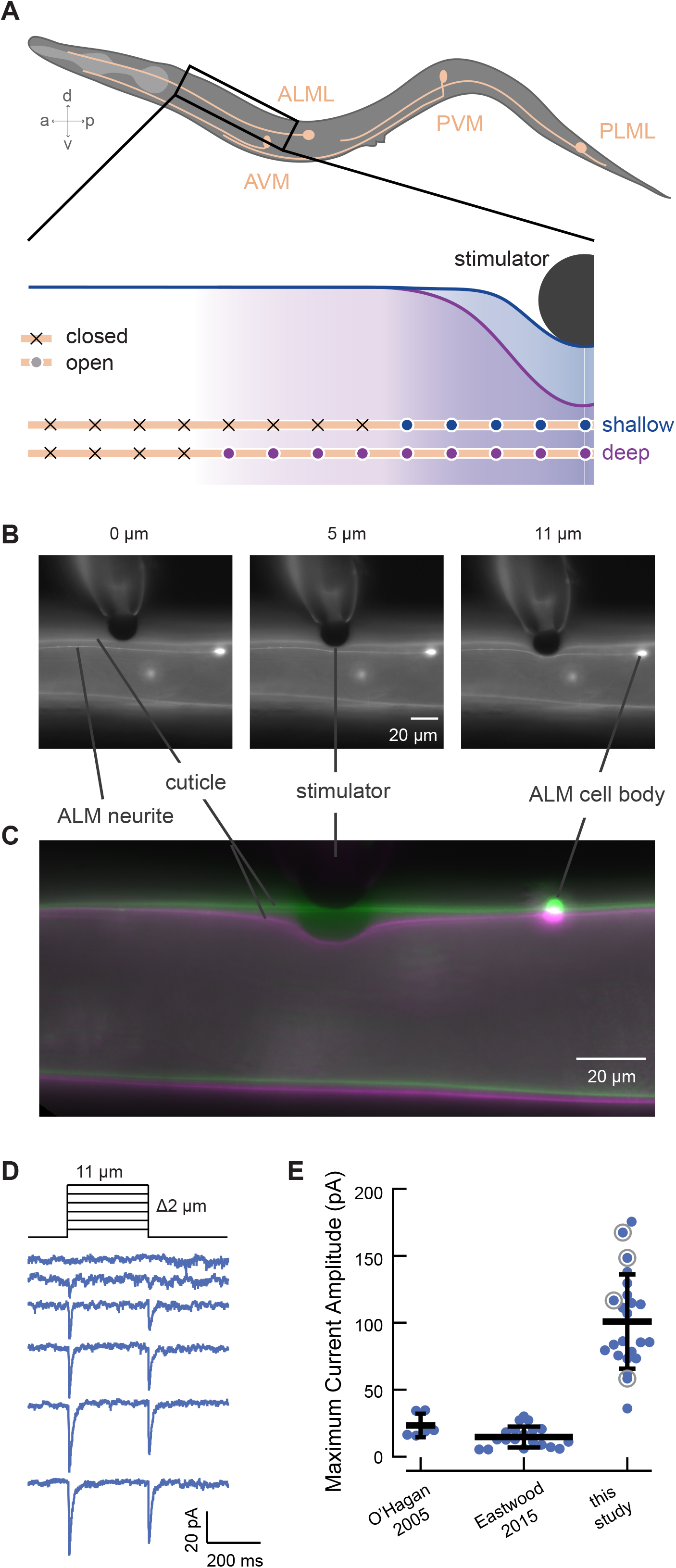
Increasing indentation increases the extent of cuticle deformation. **(A)** Illustration of the *C. elegans* six touch receptor neurons (top) and the hypothesized mechanism of channel recruitment in the ALM neuron during touch (bottom, not to scale). The channels activated by touch stimuli generating shallow (blue) and deep (purple) skin indentation are indicated by filled circles and channels that remain outside the touch-evoked strain field are indicated by ‘x’ symbols. Note that while only the channels anterior to the stimulator are shown, channel recruitment is assumed to extend symmetrically anteriorly and posteriorly. **(B)** Micrographs of stimulation of a living adult worm by a black polyethylene bead (*d* = 22 µm) at different depths. Deformation is visible in the cuticle and the ALM neuron, which are both labeled with GFP in transgenic GN865 *uIs31[mec-17p::GFP]; kaIs12[col-19::GFP]* animals. **(C)** An overlay of cuticle deformation in another GN865 animal before (green) and during (magenta) an 11-µm touch displacement with a polyethylene bead (*d* = 22µm). In these images, the ALM cell body is outside the plane of focus. **(D)** Representative mechanoreceptor currents (MRCs) activated by the application and removal of rapid displacement steps between 1 and 11µm (in 2µm increments). **(E)** Comparison of peak MRC amplitudes in two prior studies and this study. The data drawn from O’Hagan, *et al*. (2005) are responses to steps (*F* > 1µN) recorded from PLM neurons; data from Eastwood, *et al*. (2015) are responses to force-clamped steps (*F* = 1.7µN) recorded from ALM neurons. For the present study, data are drawn from displacement steps (*z =* 10µm, 25 mm/s, delivered an average of 114 ± 41µm (mean ± SD, range: 53-192 µm) from the cell body). Horizontal and vertical bars (black) are the mean and SD of each data set, respectively. Gray circles indicate values drawn from recordings shown in more detail in Figs. 3A, 3B.

### MRC amplitude depends on stimulus probe size, displacement and speed

To enable a thorough investigation of how indentation depth and speed influence activation of MeT channels in their native environment, we built a high-speed mechanical stimulator (Materials and Methods). While recording MRCs, we positioned this stimulator perpendicular to the worm’s body and used the stimulator to push a stiff glass probe carrying a large (∼22 μm diameter) polyethylene bead into living, wild-type worms (Fig. 1B and 1C). We applied displacements up to 12 µm, which is about 30% of the average body width (39 ± 4 µm, mean ± SD, *N* = 53). Unexpectedly, the peak MRCs we report here are about five-fold larger than those previously reported by us (Fig. 1E). Here, we recorded from ALM, rather than PLM. We pushed the bead perpendicularly, rather than tangentially into the worm (Árnadóttir et al., 2011; Chen and Chalfie, 2015; Eastwood et al., 2015; O’Hagan et al., 2005) and used larger beads to deliver the mechanical stimulus (Eastwood et al., 2015). We also used high-speed stimulation with minimal resonant vibration.

Intrinsically larger responses in ALM versus PLM might account for some of the difference (Chen and Chalfie, 2015), but ALM currents measured with a force-clamped stimulus (Eastwood et al., 2015) were still significantly smaller than what we measured here (Fig. 1E). Thus, the discrepancy in peak MRC size is likely to reflect differences in the stimulus paradigm or worm preparation between studies: the use of a larger bead to contact the animal’s skin, high-speed stimulation, variations in body mechanics, or a combination of these factors. For a given indentation, our model (Sanzeni et al., 2018) predicts that larger beads and stiffer animals will generate larger strain fields, which would in turn increase the number of channels available for activation. In addition to these factors, the model predicts that current amplitude increases with stimulation speed (Sanzeni et al., 2018). The indentation steps we applied here are at least an order of magnitude faster than those of the FALCON system (Eastwood et al., 2015; Sanzeni et al., 2018). Although dissection of cell bodies for recording creates variation in body stiffness that affects peak current amplitude (Sanzeni et al., 2018), this is unlikely to account for the ten-fold change in mean MRC amplitude (Fig. 1E).

Another factor affecting MRC amplitude is the expression of MEC-4, a key pore-forming subunit of the native MeT channel (O’Hagan et al., 2005). Whether visualized by immunofluorescence labeling of fixed samples (Chen and Chalfie, 2015; Cueva et al., 2007) or by direct observation of transgenic animals expressing a fluorescent protein fused to MEC-4 (Árnadóttir et al., 2011; Chelur et al., 2002; Emtage et al., 2004; Petzold et al., 2013; Vásquez et al., 2014), MEC-4 localizes to discrete puncta along TRN neurites. In prior studies (Árnadóttir et al., 2011; Chen and Chalfie, 2015; Cueva et al., 2007; Petzold et al., 2013; Vásquez et al., 2014), adjacent puncta were separated by between 1.3 and 3.8 µm and these average values were obtained by analyzing arbitrary neurite fragments. To gain a comprehensive and global view of the distribution of MEC-4 protein, we tagged MEC-4 with mNeonGreen (mNG), an exceptionally bright fluorescent protein suitable for *in vivo* applications (Hostettler et al., 2017) and developed an image processing workflow to analyze inter-punctum intervals in full-length TRNs (Materials and Methods). As shown in Figure 2, the distance between neighboring mNG::MEC-4 puncta increases with distance from the ALM cell body.

**Figure 2.**
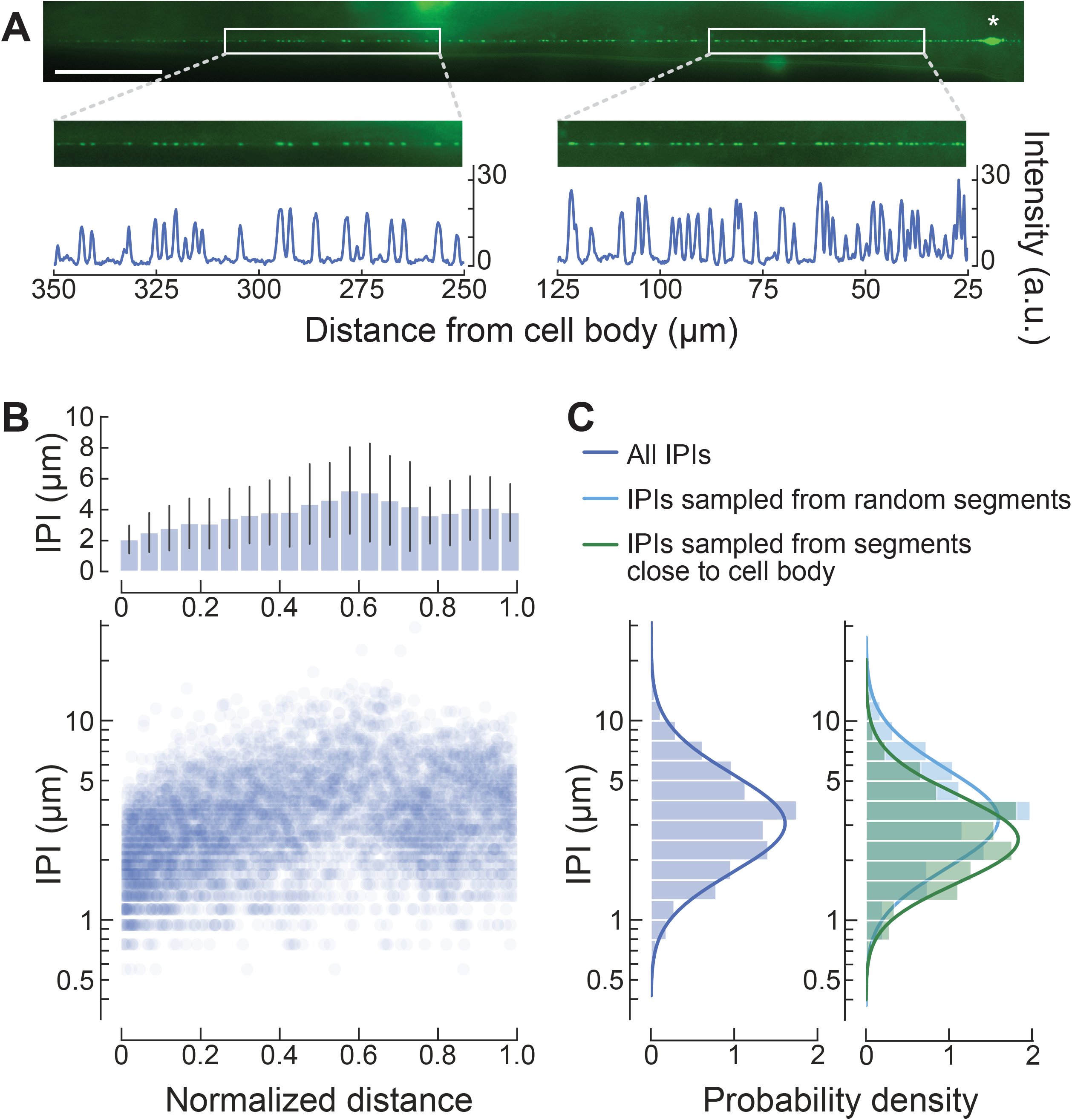
The density of MEC-4 channel puncta decreases with distance from the cell body. **(A)** mNeonGreen-tagged MEC-4 proteins localize to puncta along the entire sensory neurite. White boxes show segments distal (left) and proximal (right) to the ALMR cell body (asterisk). Fluorescence intensity profiles of the segments show a lower density of puncta (peaks) in the distal segment than in the proximal segment. Similar images were obtained from a total *N =* 51 ALM neurons; anterior is to the left; scale bar = 50 μm. **(B)** Inter-punctum interval (IPI) increases with distance from the cell body. Average IPI (± SD, top) and individual IPI values (bottom) as a function of normalized distance from the ALM cell body, a total of *n* = 5783 IPIs measured from *N* = 51 ALM neurons. **(C)** Comparison of the distribution of global IPI values (left; mean = 3.62 μm, mode or peak = 2.95 μm, SD = 2.20 μm) and IPI values derived from random (right, light blue; mean = 3.85 μm, mode = 3.24 μm, SD = 2.33 μm) or distance-constrained (right, green; mean = 2.95 μm, peak = 2.43 μm, SD = 1.60 μm) samples of 100-µm segments. The global and randomly sampled distributions are similar, while the distance-constrained condition that selected segments centered between 50-100 μm from the cell body yields a narrower distribution with a smaller average IPI.

**Figure 3.**
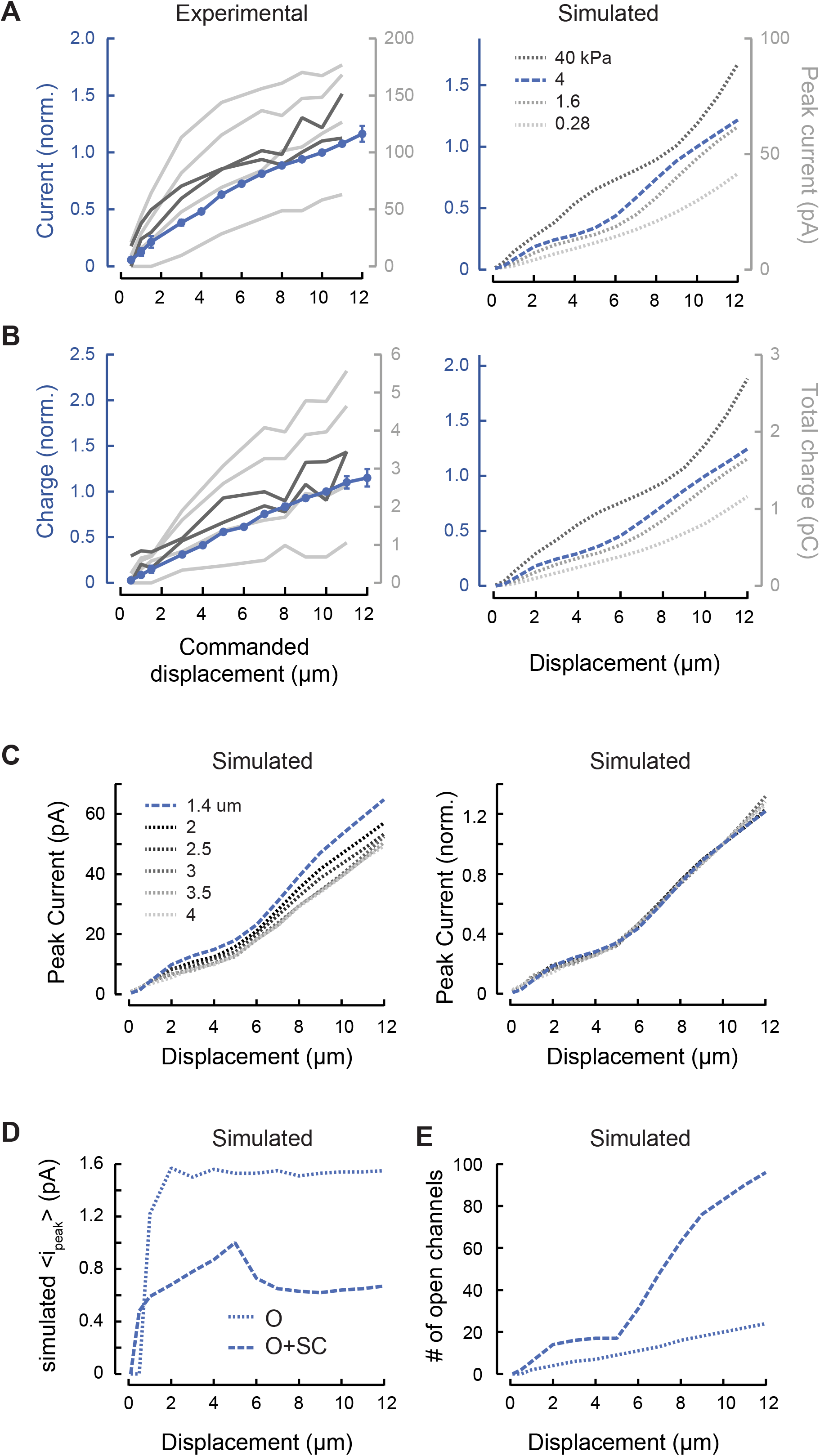
MRC sensitivity depends on body mechanics and channel recruitment. **(A)** Experimental (left) and simulated (right) peak MRC current as a function of bead displacement. Average experimental responses (blue) were normalized to the peak current evoked by a 10-µm displacement. Points are mean (± SEM) of *N* = 22 recordings. Variations among experimental dissections are evident in four individual curves (light gray) obtained following the standard dissection technique and two individual curves (dark gray) obtained from minimally dissected animals. The stimulating bead was held against the body for 250 or 300ms for displacements of 0.5-12 µm and speeds of 1.25-30mm/s. Simulations were conducted across a range of internal pressure values (mimicking variation in experimental dissections) and to identify a value (4 kPa) approximating the average experimental response. **(B)** Experimental (left) and simulated (right) total charge as a function of bead displacement, derived from the same experimental and simulated recordings as in panel A. **(C)** Increasing the simulated inter-punctum interval from 1.4μm (blue, dashed) to 4 μm (dotted) decreases peak current (left), but does not affect the shape of normalized current-displacement curves (right). Simulation parameters for each IPI are in Table 1. **(D)** The simulated mean peak current per activated channel increases with small displacements but saturates at larger displacements. A threshold of 0.5 ∗ *i*_*o*_ reflects the open probability of channels in the open (O) state (solid line) while a threshold of 0.1 ∗ *i*_*o*_ includes both O and sub-conducting (SC) channels (dotted line). **(E)** The number of simulated channels in the O or SC state increases continuously with displacement and does not appear to saturate. Calculated using the same thresholds as used in panel C.

Pooling inter-punctum intervals without regard to position along the neurite yields a log-normal distribution (Fig. 2C), reminiscent of the distribution first reported by Cueva *et al*. (2007). This finding suggests an explanation for the variation in prior reports of average inter-punctum intervals—namely, that datasets sampling fragments closer to the cell body would yield smaller inter-punctum intervals than those sampling fragments further away. We tested this idea computationally, by sampling 100µm fragments centered at distances of 50 to 100µm from the ALM cell body, matching the delivery of mechanical stimuli in this study (Fig. 2D, green). As expected, this resulted in a narrower distribution with a shorter average inter-punctum interval than when we sampled 100μm fragments at any distance (Fig. 2D, blue). Thus, mechanical stimuli delivered near these positions might be expected to evoke larger MRCs than those delivered further away, as seen in the PLM neuron (Chen and Chalfie, 2015). This factor is unlikely to account for the ten-fold change in mean MRC amplitude (Fig. 1E), however, since the present and prior (Eastwood et al., 2015; O’Hagan et al., 2005) studies also relied on stimuli delivered approximately 100µm away from the cell body. Based on these considerations, we infer that the probe size and stimulus speed are the dominant factors that account for the larger currents.

### Peak MRCs increase with stimulus size, but do not saturate

Indentation depth also affects peak current amplitude and total charge transferred (Fig. 3A and 3B, left). Increasing the indentation depth has two effects, as demonstrated in our computational model linking tissue mechanics to MeT-channel activation in living animals (Sanzeni et al., 2018). The first effect is an expansion of the strain field, which increases the number of channels available for activation. The second is an increase in the probability that a given MeT channel is activated. This model also shows that the channels closest to the point of stimulation, where the deformation is greatest, are more likely to open than distant channels (Fig. S1). These two effects combine to determine MRC peak amplitude as well as the total charge transferred and result in non-linear, non-saturating current-indentation curves (Fig. 3A and 3B, right).

Although we expect variations in body stiffness and internal pressure due to natural variation and variation in the dissection procedure (Eastwood et al., 2015), these parameters are largely outside of experimental control. To circumvent this technical limitation, we used simulations based on the model of Sanzeni *et al*. (2018) to systematically explore the influence of variations in pressure (Fig. 3A and 3B, right). With the exception of internal pressure, the free parameters (Table 1) of the simulation were derived by fitting a dataset collected with a different stimulation system (Eastwood et al., 2015). Higher internal pressures correspond to stiffer animals and lower pressures correspond to softer ones. The simulated current amplitude for a 10-μm indentation is higher in stiff animals compared to softer ones (Fig. 3A). Henceforth, the simulations rely on the parameters in Table 1 and an internal pressure of 4 kPa, which reflects an intermediate stiffness.

To test the effect of body stiffness *in vivo*, we obtained two high-quality recordings under the minimally dissected (‘stiff’) regime described in Eastwood *et al*. (2015). While these currents were larger than the average, they fell within the observed range for dissected preps in this study (Fig. 3A-B, dark grey). This may be due to inadvertent selection wherein we were only able to successfully isolate cell bodies from the least pressurized animals, but we cannot distinguish between these possibilities in the present data set. Alternately, the factors that we believe led to larger currents in this study (larger bead size and faster stimulation) may have minimized the effect of variations in stiffness.

In simulations, we used a fixed inter-punctum interval of 1.4µm, a value drawn from immunofluorescence labeling of native protein (Cueva et al., 2007). To determine how this choice affected our results, we repeated simulations with values between 1.4 and 4 µm. For each value, we determined single-channel model parameters from the best fits to recorded neural responses as discussed in the Materials and Methods section (Table 1). We found that the peak current decreased slightly with the inter-punctum interval (Fig. 3C, left), but that the shape of the relationship between displacement and normalized peak current was unaffected (Fig. 3C, right). These simulations show that, while the quantitative aspect of our results depend on the inter-punctum interval chosen, the scaling relations found are robust to the specific choice. In the following plots, we report simulations performed with an inter-punctum interval of 1.4µm; by using the smallest value within the physiologically reported range, we avoided position-related artifacts in the responses to small displacements.

In all prior studies of MRCs in *C. elegans* touch receptor neurons, current-indentation curves for MeT channels have been treated as a saturating function of stimulus size. These experimental and computational results indicate that such a treatment neglects the influence of tissue mechanics as well as the de-localized nature of MeT channels. Furthermore, our results show that the shape of the current-indentation curve is independent of interpunctum interval but varies with other factors, such as body stiffness, that are extrinsic to the MeT channels themselves.

Using the Sanzeni *et al*. (2018) model, we further examined the effect of displacement on current by separating the effect of indentation on open probability from the effect of indentation on increased channel recruitment via an expansion of the strain field. In this model, the strength of the stimulus received by each channel and, hence, its predicted open probability depends on its location relative to the stimulus. Based on single-channel recording that revealed a robust sub-conductance state in MEC-4 channels (Brown et al., 2008), we included a sub-conductance state between closed and open states in the model, and derived the sub-conductance current *i*_s_ relative to the open-channel current *i*_o_ from the fit (Table 1). Sub-conductance and fully-open states carry 0.35 and 1.6 pA of inward current, respectively with the primary parameter set. (Subconductance current ranged from 0.35 pA to 0.84 pA when fit with different IPIs (Table 1)). These values are consistent with both single channel recordings of channels expressed in heterologous cells (Brown et al., 2008) and estimates derived from non-stationary noise analysis (O’Hagan et al., 2005).

As a concise method of describing the single channel response, we computed the mean of the peak current carried by each channel, regardless of whether we considered only channels that reach the open state (designated ‘O’) or also included channels in a sub-conductance state (designated ‘O+SC’). Based on this measurement, we observed that the mean current per channel rose with small displacements, but then held steady (Fig. 3D). The inclusion of sub-conducting channels more closely reflects the behavior of both current and charge. As such, all subsequent plots of this type reflect the O+SC condition. In contrast, the number of open channels continued to increase with increased displacement, behaving similarly to the current (Fig. 3E). The discontinuity in the number of channels that occurs near 5μm displacement is due to a non-linear recruitment of a population of distant channels with low open probability (Fig. S1), which in turn decreases the average current per channel (Fig. 3D). Collectively, these results suggest that channel recruitment dominates the overall current-indentation relationship, while increases in open probability and transitions from sub-conducting to open states steepen the curve, especially for small indentations.

### MeT channels are activated by distant stimuli

Our results show that indentation at one point affects multiple channels along the neurite. Thus, we expect that MRC amplitude depends on the overlap between the strain field and the neurite (Fig. 4A). We explored this concept experimentally by holding the stimulus size constant and varying the distance between the ALM cell body and the stimulator in both the anterior and posterior directions (Fig. 4B). For recordings obtained while stimulating anterior to the cell body, we corrected for errors arising from the fact that the TRNs are not isopotential (O’Hagan et al., 2005; Materials and Methods). Although we did not detect a systematic relationship between current amplitude and distance from the cell body in the anterior direction (Fig. 4B, blue), the high variation in current amplitudes across individual recordings could mask such a relationship. Additional experiments involving measurements at several positions during a single recording may be required to resolve this uncertainty.

**Figure 4.**
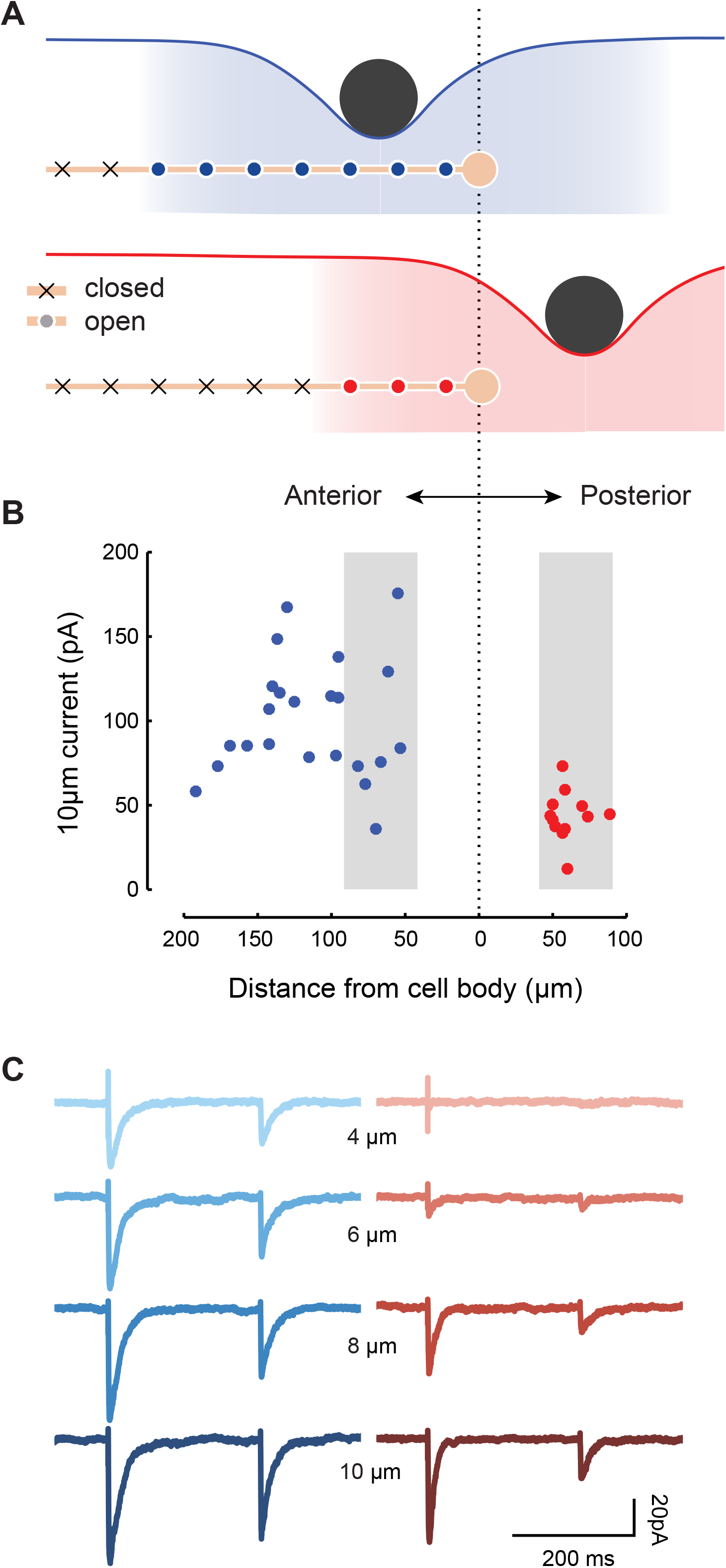
MRCs are evoked during stimulation at a distance from the nearest channels. **(A)** Diagram illustrating how positioning a stimulus anterior to the ALM cell body (top) is predicted to activate more channels (blue or red circles) than positioning the same stimulus posterior to the cell body (bottom). **(B)** Peak MRCs evoked by a 10-μm step (duration: 250ms; speed: 25 mm/s) are generally larger when the stimulator is positioned anterior to the ALM cell body (blue) than when it is posterior (red). Each dot is a single recording; for anterior recordings (blue), MRC size was for space-clamp error (see Methods). Gray shading indicates recordings evoked by stimuli delivered 40-90 μm away from the cell body. **(C)** Representative MRCs for an anterior stimulus (left, blue traces, *n* = 3 technical replicates at 53 μm distance) or posterior stimulus (right, red traces, *n* = 4-5 technical replicates at 58 μm distance). Similar results were obtained from a total of *N* = 22 and 12 recordings evoked by stimulating anterior and posterior to the cell body, respectively. For all recordings, the stimulating bead was held against the worm’s body for 250 or 300 ms for displacements of 0.5-12 µm and speeds of 1.25-30 mm/s.

There are no channels from ALM posterior to its cell body. Despite this fact, stimuli delivered posterior to ALM can evoke calcium transients in the ALM cell body (Suzuki et al., 2003), which are likely driven by MRCs (Fig. 4B). On average, these currents had smaller amplitudes than currents elicited by stimuli of an equivalent magnitude anterior to the cell body (Fig. 4B and 3C). Although the variation in current amplitude was lower than the variation we observed for anterior stimulation, there was no obvious relationship between current amplitude and distance within 100 µm of the cell body in the posterior direction. For more distant stimuli, however, we could not detect currents even for large displacements (*N* = 4).

Consistent with the reduced overlap between the strain field and the neurite for posterior stimulation, larger displacements were necessary for eliciting currents under this condition (Figs. 4C, 5A, 5B). Peak current-displacement curves were shifted to the right relative to curves observed for anterior stimulation such that the posterior stimulation only evoked consistently detectable MRCs for indentations larger than 4 μm. Simulations captured this shift in peak current-indentation curves (Fig. 5A, right) and suggested that the size of the shift is proportional to the distance from the cell body. We also plotted the total charge—the area under the MRC curve—against displacement to account for differences in the timing of channel opening. This reduced the difference in operating range in our experimental data (Fig. 5B, left), but not in the simulations (Fig. 5B, right). This suggests that channels activated during posterior stimulation are activated in a less synchronous manner than channels activated during anterior stimulation. For both measures, larger stimuli are necessary for activating channels in the simulations.

**Figure 5.**
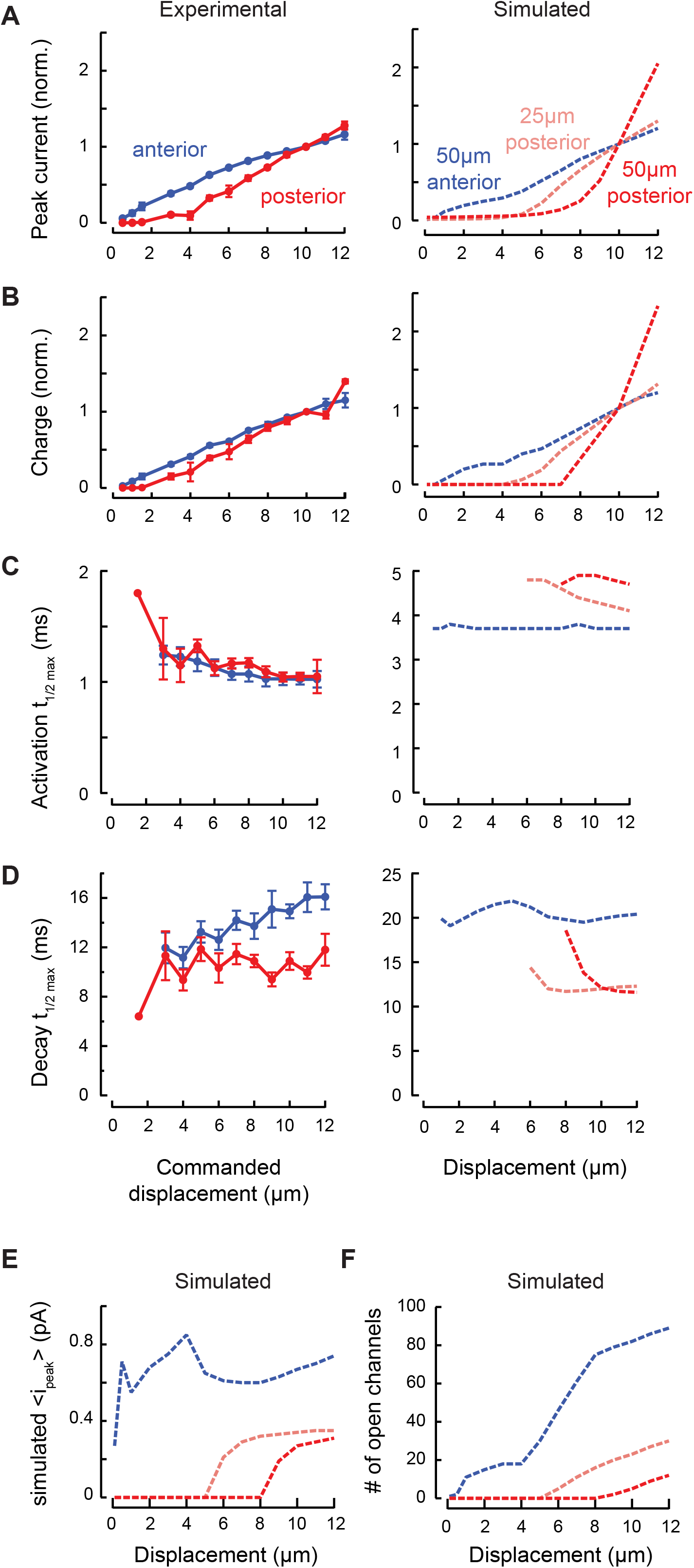
MRC sensitivity depends on stimulus position. **(A)** Experimental (left) and simulated (right) normalized peak MRC current-displacement curves in response to stimuli delivered anterior (blue) and posterior (red) to the cell body. Experimental points are mean ± SEM for *N* = 22 worms with anterior stimulation (blue, same recordings as in Fig. 3A) and *N* = 12 worms with posterior stimulation (red, stimulator positioned a mean ± SD of 61 ± 12µm posterior to the ALM cell body with range 48 – 89 μm, recordings indicated in Fig. 4B). Simulated traces (right) show predicted normalized peak currents for stimuli delivered anterior (blue, 50 µm from cell body) and posterior (pink, 25 μm; red, 50 μm) to cell body. **(B)** Experimental (left) and simulated (right) relationship between total MRC charge and displacement, derived from the experimental and simulated data in A. **(C)** The time to reach half-maximal current is similar in anterior and posterior (left) recordings, but diverges in simulations (right). **(D)** The decay half-time increases with displacement for anterior, but not posterior recordings (left), a distinction that is not reproduced in simulations (right). Experimental results in panel C and D were limited to recordings obtained a stimulator positioned within 90 μm of the cell body (indicated in Fig. 4B, gray) and include a total of *N* = 7 (anterior) and *N* = 11 (posterior) recordings. **(E)** The simulated mean peak current per active channel is lower for stimuli delivered posterior to the cell body than it is for the anterior stimuli. **(F)** Simulations reveal that stimuli recruit channels only if the displacement is large. The threshold of 0.1 ∗ *i*_*o*_ for active channels includes both open and sub-conducting channels.

Given the discrepancy between peak current and total charge, we looked at how the activation and decay rates varied with displacement in anterior and posterior recordings. Passive conduction over a long distance can broaden the time course of electrical events measured at the cell body (Bekkers and Stevens, 1996; Rall, 1967). Therefore, we limited this analysis to stimuli within 90 μm of the cell body—less than the average length constant λ = 96 ± 3 μm (mean ± SEM, *N* = 83) calculated for our recordings—for both anterior and posterior recordings.

We estimated activation rates by measuring the time required for the current to rise to half the peak value from the start of the stimulus, and decay rates from the time to peak to when the current dropped to half the peak value. Activation rate increased with stimulus size and was similar for anterior and posterior stimulation (Fig. 5C, left). We failed to detect any difference in latency. The decay rate for anterior—but not posterior—stimulation increased slowly with displacement (Fig. 5D, left). This observation is consistent with space-clamp error for channels beyond the point of stimulation (the point at which we corrected for voltage attenuation): larger stimuli affect channels that are much further away from the cell body and the contribution of these channels to the total current measured at the cell body is slowed and decreased by passive conduction. Indeed, when we included recordings in which the stimulator was placed even further anterior to the cell body, we found that both the activation and decay were slower (data not shown). Channels activated by posterior stimulation are located near the cell body, where this error is low, but their position at the edge of the strain field means they are activated only weakly. The simulation does not consider space clamp error, but predicts faster decay with posterior stimulation because tissue experiencing low strain only needs to move a short distance to relax. Although the smaller shift of total charge is not entirely explained by this finding, simulations of activation and decay rates agree qualitatively with the experimental results in at least one other way: the further posterior the stimulator is placed, the larger the displacement required to reach any channels in the neurite (Fig. 5C and 5D, right). The simulation predicts slower activation times for posterior stimulation because strain takes longer to reach distant channels than proximal channels. However, we could not clearly detect such a difference in our experiments with this fast step protocol.

As noted above, the macroscopic currents we record depend on 1) stimulus-dependent changes in a channel’s open probability; and 2) the number of channels exposed to each stimulus. Using average peak single-channel current as a proxy for open probability, our simulations predict that channel open probability saturates at smaller displacements for anterior stimuli than for posterior stimuli (Fig. 5E). Also, the fact that larger stimuli reach more channels than smaller stimuli is the dominant factor controlling peak current-displacement curves for both anterior and posterior stimuli (Fig. 5A).

### MRC size and activation rate depend on stimulus speed and direction

The peak amplitude of MRCs in *C. elegans* TRNs increases with the speed of indentation (Fig. 6A, 7A left). This could reflect speed-dependent activation of individual channels. It could also reflect a speed-dependent increase in synchrony, or in-phase activation, of a population of channels. To differentiate between these possibilities, we analyzed peak current and charge across a wide range of speeds and with fine resolution. Unlike peak MRC size, charge accounts for all channel openings regardless of timing. It follows, therefore, that if the speed-dependence of peak MRC amplitude were entirely due to variations in the phase of channel opening, then the charge would be independent of speed. Contrary to this prediction, we found that charge was speed dependent in both experiments and simulations (Fig. 7B). Collectively, these findings indicate that variations in the phase of channel activation are not sufficient to account for the speed dependence of MRC generation.

**Figure 6.**
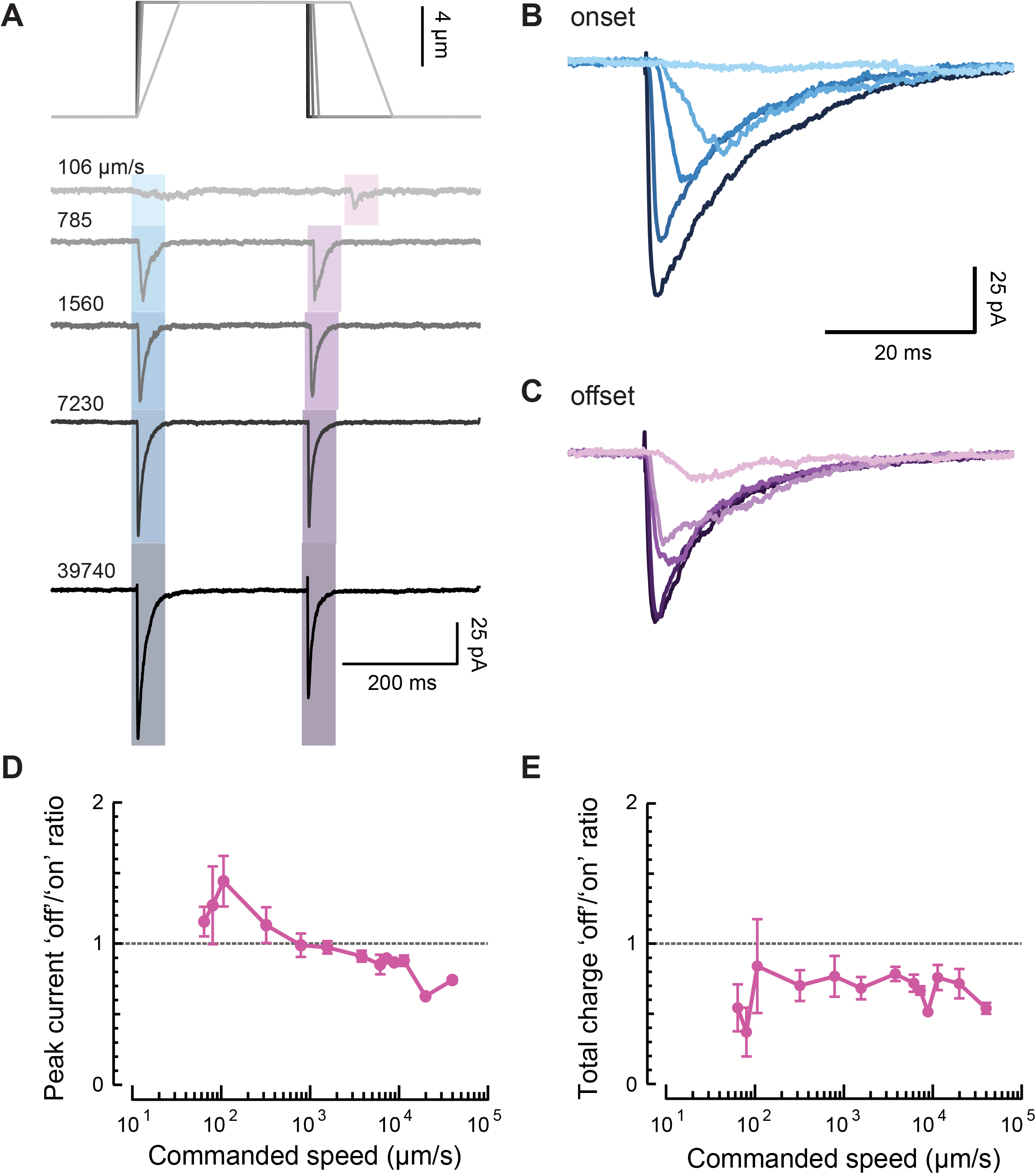
Stimulus speed increases MRC amplitude and hastens activation. **(A)** Representative recording (average of *n* = 2-7 technical replicates) of MRCs evoked by trapezoidal stimulus patterns (constant displacement, variable onset and offset speed). **(B, C)** Expansion of MRCs at the onset **(B)** and offset **(C)** of a trapezoidal stimulus. Currents at the onset are larger at fast speeds, but smaller and slower to activate at low speeds. **(D, E)** Ratio of ‘off’ response relative to ‘on’ response amplitudes, measured as either peak current **(D)** or total charge **(E)** for *N* = 6 worms stimulated at an average of 67 ± 10 μm (mean ± SD) anterior to the cell body.

**Figure 7.**
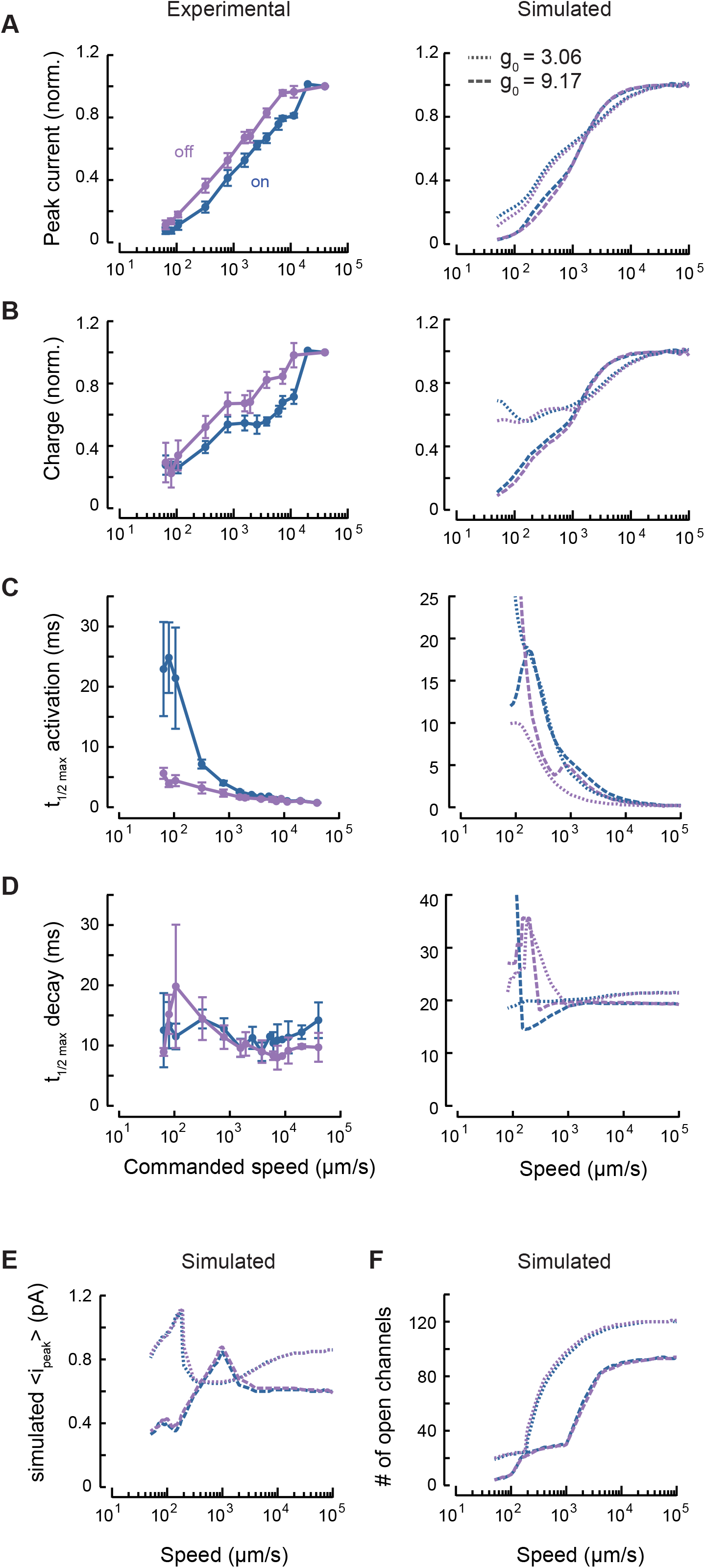
Synchrony in channel opening partially explains ‘on’/’off’ asymmetry. **(A)** Experimental (left) and simulated (right) peak current as a function of stimulus speed, normalized to the response a stimulus speed of 40 mm/s. Responses to stimulus onset (‘on’, blue) and offset (‘off’, purple) are plotted separately. Points are mean ± SEM (*N =* 6 recordings). Simulations were performed with the default force required to activate a channel (dashed, *g*_0_ = 9.17) or a smaller activation force (dotted, *g*_0_ = 3.06). **(B)** Experimental (left) and simulated (right) total charge as a function of stimulus speed, derived from the same experimental and simulated recordings as in panel A. **(C)** Experimental (left) and simulated (right) ‘on’ currents activate more slowly than ‘off’ currents at slow speeds. **(D)** The MRC decay rate is similar for both ‘on’ and ‘off’ currents; simulated data (right) show similar behavior at high speeds. For the data in panels A-D, the stimulus bead was held at a constant displacement of 8µm in between the onset and offset ramp delivered at the indicated speeds and positioned 67 ± 10 μm (mean ± SD) anterior to the cell body, on average. **(E)** Mean peak single-channel current from the simulation increases with speed and then saturates. The threshold of 0.1 ∗ *i*_*o*_ for active channels includes both open and sub-conducting channels. **(F)** The number of open channels increases with speed, but saturates at high speeds. The same thresholds are used as in E.

In contrast to displacement sensitivity, measured current and charge saturate at high speed and simulated current and charge clearly saturate at speeds above 1,000 μm/s (Fig. 7A, 7B left). This finding suggests that indentation depth governs the number of channels eligible for recruitment. To estimate the relative contributions of channel recruitment and gating to speed dependence, we turned to simulations, which predicted a similar dependence on speed as with displacement (Fig. 7A-D right). We found that both peak channel current and the number of open channels saturate at high speeds (Fig. 7E, 7F) and that the number of open channels increases with stimulus speed (Fig. 7F).

The saturating number of channels depends on the value of *g*_0_, a model parameter controlling the effect of linker elongation on channel opening (Fig. 7F). Smaller parameter values render the channels more sensitive to linker elongation, allowing recruitment of more channels. Thus, for a fixed final displacement there seems to be a maximal number of channels that can be opened and this number is expected to be inversely proportional to the force needed to activate single channels. Even with *g*_0_ and the final displacement held constant, channel recruitment increases with stimulus speed; this results from the larger forces generated by quicker changes in strain. Thus, MRC amplitude increases with speed because more distant channels are activated in response to faster stimuli.

Both ‘on’ and ‘off’ mechanoreceptor currents increase in size with stimulus speed (Figs. 6A-C and 7A and 7B, left), but to different extents. At low speeds, ‘off’ MRCs have a larger peak current than ‘on’ MRCs, but the ratio reverses at high speeds (Fig. 6D). This observation was not recapitulated in simulations, suggesting that its origin lies in an element that is absent from the present computational model or an assumption that may not hold. For instance, the model assumes that tissue indentation follows the stimulator speed precisely. It is possible, however, that the stimulator bead detaches from the body during fast ‘off’ stimuli creating asymmetries in the mechanical loads delivered at high speed. Above, we argued that synchrony or phase differences in channel activation do not fully account for speed-dependence generally. When we explored the possibility that phase differences contributed to the ‘on’/‘off’ asymmetry observed in peak current amplitudes by plotting the ratio of ‘off’ to ‘on’ charge against stimulus speed, we found that ‘off’ responses consistently carried less charge than ‘on’ responses. This ratio was not obviously speed dependent (Fig. 6E).

‘On’ currents activate slowly at slow speeds, while ‘off’ currents activate rapidly and vary less with stimulus speed in both experimental and simulated data (Fig. 7C). The simulation predicts that recruiting more channels by decreasing *g*_0_ would increase the asymmetry in activation rate (Fig. 7C, right). This is not true for the decay rate, however. Both ‘on’ and ‘off’ MRCs seem to decay at a relatively constant rate regardless of stimulus speed (Fig. 7D). This suggests that activation depends on stimulus speed, while decay depends on tissue relaxation. Intuitively, we might expect channels to be more likely to be opened in-phase during ‘off’ stimuli because all channels affected by the deformation will start moving once the stimulator moves. In contrast, during slow ‘on’ stimuli, channels closer to the stimulator will be affected before the deformation reaches further channels, resulting in out-of-phase opening and slower MRC activation. This is supported by the observation that activation of ‘on’ MRCs appears to be faster at high speeds (Figs. 6B and 7C). The activation rate of ‘off’ currents is much less dependent on stimulus speed (Figs. 6C and 7C), but the amplitude of ‘off’ currents still increases with speed despite this greater synchrony (Fig. 7A), reinforcing the idea of another mechanism for speed dependence.

### Sinusoidal stimuli evoke steady MRCs in a frequency-dependent manner

The stimuli a worm encounters in the wild or in the laboratory (Nekimken et al., 2017b) involve complex signals with a variety of frequencies. Prior work shows that the TRNs seem poised to respond to variable signals—calcium responses to sinusoidal “buzz” stimuli are much stronger than those evoked by simple steps (Fehlauer et al., 2018; Nekimken et al., 2017a; Suzuki et al., 2003). Therefore, we systematically examined the frequency response to sinusoidal mechanical stimuli at frequencies ranging from 10 Hz to 500 Hz. The representative traces in Fig. 8A show both the signal from the photodiode measuring bead motion as well as the currents evoked by this stimulation. We see little evidence of adaptation following the application of a buzz-like stimulus: MRCs evoked by a step stimulus before and after a 1 s-sinusoid show no more than 10-15% attenuation. This result starkly contrasts with studies on Piezo1 and Piezo2 channels, which adapt following sinusoidal stimulation (Lewis et al., 2017).

**Figure 8.**
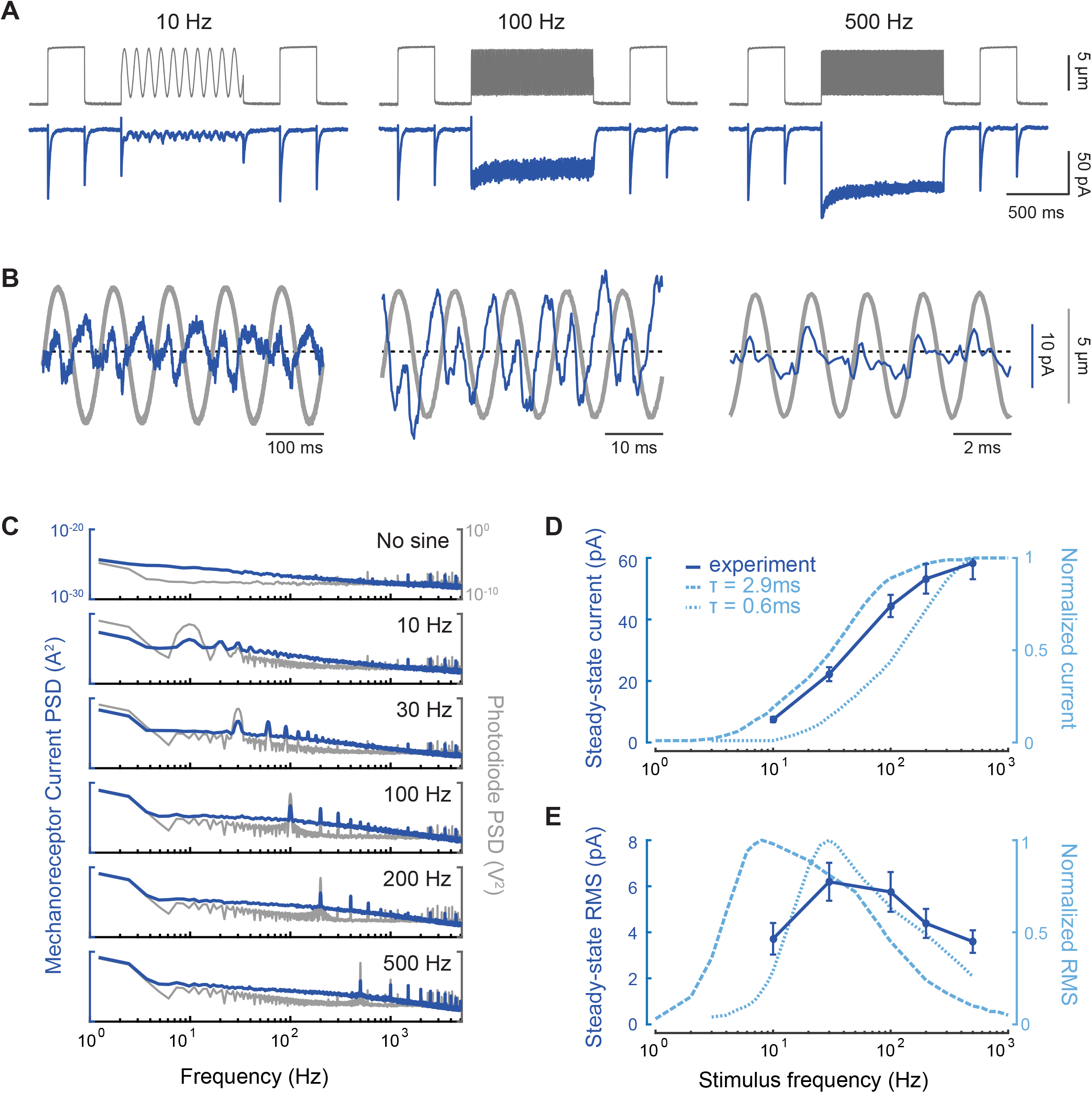
MRCs evoked by sinusoidal stimuli have complex, non-linear temporal dynamics. **(A)** Representative traces of MRCs (blue) evoked by sinusoidal stimuli (mean displacement: 5 µm; amplitude: 8 µm) delivered anterior to the cell body. Gray traces are photodiode-based tracking of stimulator bead motion. Flanking displacement pulses demonstrate recording quality and stability. Each trace is a mean of *n =* 5-6 trials; similar results were obtained in *N =* 7 recordings. **(B)** Expanded responses from panel A (blue) overlaid on the photodiode measurement of bead motion (gray). The dashed line indicates the mean current. **(C)** Comparison of the power spectrum of MRCs (blue) and bead motion (gray). The power at the first and second harmonics is similar for MRCs, but not for bead motion. Traces are the average from *N =* 7 recordings. **(D)** Steady-state mean current increases as a function of stimulus frequency. The experimental current-frequency curve (dark blue, solid) is compared to simulations (light blue, dashed) using two values for the relaxation time, τ, of the hypothetical elastic filament in the model. **(E)** Steady-state RMS current shows a band-pass frequency dependence. Comparing experimental (dark blue, solid) and simulated results (light blue, dashed) demonstrate that frequency dependence depends on the value of the relaxation time, τ. Faster relaxation times shift activation to higher frequency. Points in panel D and E are the mean current (± SEM) and mean RMS (± SEM), respectively, derived from the current measured during the last 100 ms of each sinusoidal epoch in 7 recordings. The stimulator was placed 72 ± 13 μm (mean ± SD) anterior to the cell body.

In agreement with our previously reported experimental and simulation results (Eastwood et al., 2015; Sanzeni et al., 2018), the expanded traces show that inward currents occur at twice the frequency of the stimulus (Fig. 8B), especially at frequencies below 500 Hz. This suggests an alternation between ‘on’ and ‘off’ responses. We next examined the power spectral density of the stimulus and the MRC response. We found that MRCs have roughly equal power at 1x and 2x the stimulus frequency (Fig. 8C), whereas the spectrum of the stimulator shows a larger peak at the stimulus frequency and a smaller one at its second harmonic. The relative power of the second harmonic peak in the MRC response decreases slightly at higher frequencies, which is reflected in the decreasing asymmetry between subsequent (’on’ vs. ‘off’) peaks (Fig. 8B). This finding demonstrates that the neuron’s response to stimulation is non-linear.

The average steady-state current continues to increase with frequency (Fig. 8A, 8D). Although the experimental data are shifted to the right relative to the predictions of the model with the previously set parameters (τ = 2.9 ms, dashed line), both approaches show the tissue acting as a high-pass filter (Fig. 8D). This is consistent with TRNs being relatively insensitive to slowly varying stimuli and thus ignoring self-motion. By contrast, the variance in the current, as measured by the root mean square (RMS) of the response at steady state, shows band-pass behavior (Fig. 8E). Again, the peak of the experimental curve is shifted to higher frequencies than the model predicts with the previous parameters (dashed line). The band-pass behavior in Fig. 8E can be understood as being due to the fact that channels do not have time to transition to lower-conductance states during high-frequency stimulation. Because the channels are held in an open or sub-conducting state, their contribution to the RMS noise declines with frequency.

We hypothesized that the simulated results were shifted to the left because channels were not able to close in phase with the sinusoidal stimulus. To test this possibility, we shortened the time required for the elastic filament to relax to baseline (τ = 0.6), which allowed us to better match the peak RMS (Fig. 8E, dotted line). However, this change also shifted the steady-state current to the right. Collectively, these observations suggest that additional modifications to model parameters will be needed to fine-tune the mechanical filter that links touch sensation to MeT channel activation.

### Single channel activation is a function of position and stimulus intensity

Our model allows us to explore spatial and temporal dynamics not only at the level of macroscopic currents, but also at the individual channel level. We selected four channels at varying distances from one realization of the simulation, and calculated the expected ‘on’ and ‘off’ responses to various step displacements, variable speed trapezoidal profiles, and sine frequencies. Our goal was to examine how MRC dynamics vary along the length of the neurite and as a function of stimulus type.

The simulated channel current reflects the weighted probability that a channel at this location is in an open or sub-conducting state. The maximal current possible is −1.6 pA, which is the value calculated from the measured single-channel conductance (Brown et al., 2007) and our recording conditions. Channels that are more distant from the center of the stimulator only respond to large displacements (Fig. 9A, highlighted). Channels directly under the stimulator (A, rightmost column) reach a fully open state even with small displacements; with larger displacements, the open probability remains high for longer periods of time even when the step takes the same amount of time. Because the model parameters were fit based on a previous dataset, the total current predicted by the simulation (Fig. 9A, left panel) is smaller than what we observed at high speeds. This discrepancy could be due to uncertainty in some assumptions used in the model. For instance, channels in the model were distributed at regular 1.4 μm spacing, which is shorter than the average 2.95 μm spacing we observed in the region we were stimulating (Fig. 2). However, increasing the inter-punctum interval would lead to smaller simulated currents. Another possibility is that individual puncta contain more than a single functional channel. Additional investigations of the response to small (1µm or less) displacements with small stimulators, super-resolution studies of channel protein, or both are needed to gain further insight into this question.

**Figure 9.**
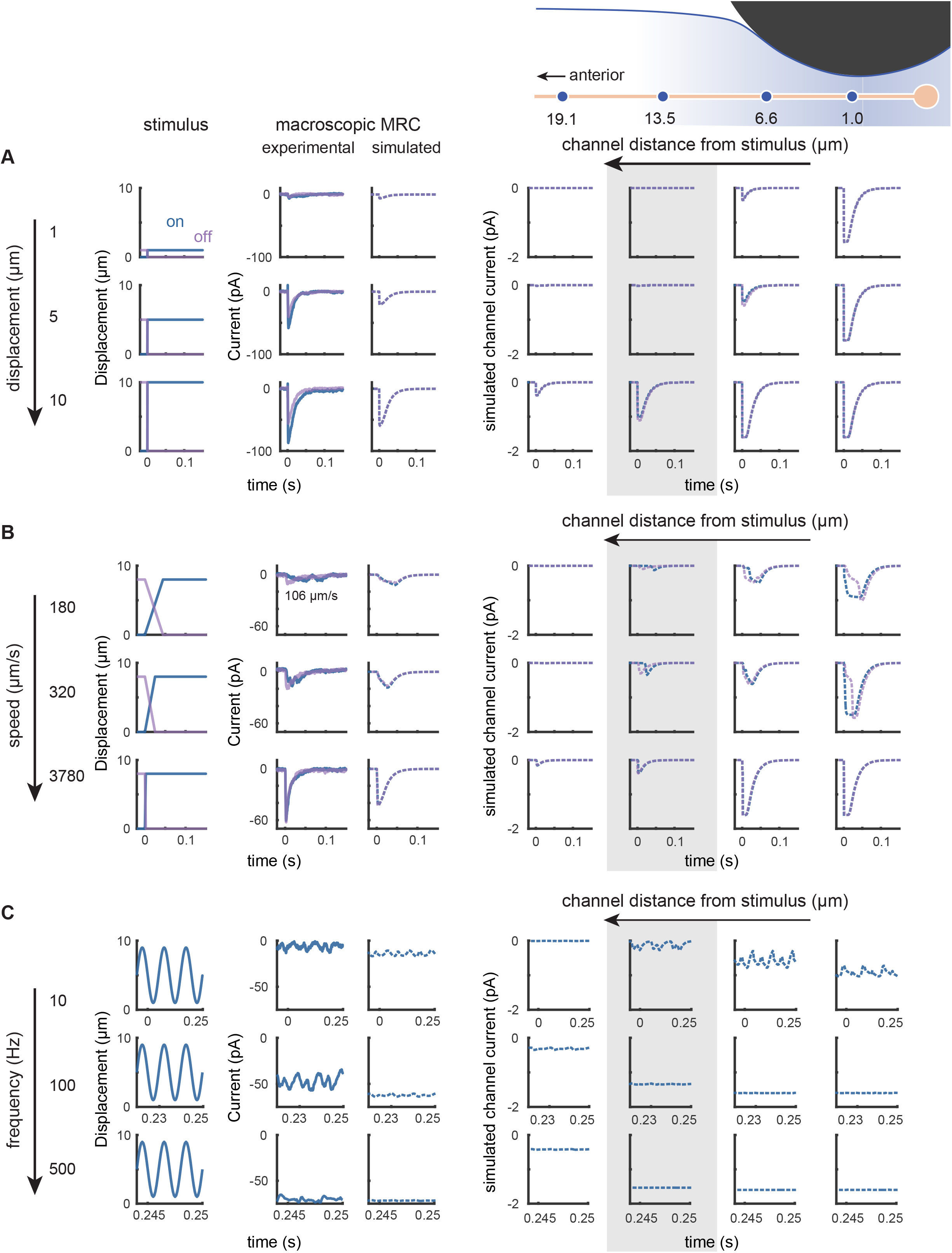
Simulations reveal that single-channel responses depend on channel position and stimulus intensity and speed. **(A)** Increasing displacement evokes larger macroscopic currents at stimulus onset (blue) and offset (purple) by recruiting distant single channels. The gray box highlights a distant channel that is unaffected by smaller displacements. **(B)** Increasing speed evokes larger macroscopic currents at stimulus onset (blue) and offset (purple) by increasing in-phase activation and open probability, and by recruiting more distant channels. ‘On’ current through the highlighted channel (gray box) has a higher latency at slow stimulus speeds, but matches the ‘off’ current at fast speeds. Experimental traces in Panel A (steps) and B (trapezoids) are from the same recording, obtained with the stimulator positioned 95µm anterior to the cell body. The slowest experimental trapezoid was delivered at 106 μm/s rather than 180 μm/s as in the simulation. **(C)** Increasing stimulus frequency evokes larger steady-state current with smaller RMS variation by recruiting more distant channels and increasing open probability. Experimental traces are from a single recording (stimulator position = 69 μm anterior) selected from the last 100-ms epoch of a 500-ms sine wave stimulus period. In panels A-C, the first three columns are stimuli at the onset (blue) and offset (purple) of each stimulus pattern, representative traces of observed currents for the stimulus in each row, and simulated currents. The last four columns are simulated currents carried by individual channels positioned either (left to right): 19.1, 13.5, 6.6, or 1 μm anterior to the stimulator. Diagram showing stimulator and channels is for visual guidance (not to scale).

When we held displacement constant and varied the speed, we found that ‘on’ and ‘off’ currents activate faster at higher speeds through a combination of synchrony and increasing open probability. Channels distant from the stimulus take longer to receive and respond to ‘on’ stimuli at slow speeds (Fig. 9B, highlighted). Channels at the point of stimulation (B, rightmost column) behave in the opposite manner: although the ‘off’ current initially rises like the ‘on’ current, it then slows and takes longer to reach the peak. Intermediate channels (B, second to right column) open both faster and more fully with increased speed. This dependence on channel distance is inherited from the underlying strain dynamics and explains the predicted slow offset activation times (Fig. 7C), which depend more strongly on the most proximal channels, and reflects the fact that the time course of strain rate depends on distance from the stimulus.

The model nicely captures the increase in steady-state current that occurs when we increase the frequency of sinusoidal stimuli. However, the fluctuations in the current (measured earlier as the RMS) are larger experimentally (Fig. 9C, left panels). In both the observed and simulated data, the relative amplitudes of the peaks at 2x the stimulus frequency also decrease with frequency. Thus, the ‘off’ response seems weaker. The open probability for channels near the stimulus remains more constant at higher frequencies (Fig. 9C, highlighted column). The intensity of the stimulus reaching the channel is attenuated in distal channels, as seen by their lack of response to smaller or slower indentations (Fig. 9A, 9B), which contributes to the decrease in RMS.

### Single channel sensitivity also determines how many channels are opened

Having established the relationship between stimulus profile and the activation of channels proximal and distal to the stimulator, we next sought to derive an estimate of the energy required to activate single channels. Following Elmi *et al*. (2017), we calculated the strain energy density (SED), a measure related to compressive strain that has been used as a proxy for the stimulus experienced by individual mechanoreceptors in various models of somatosensation (Dandekar et al., 2003; Elmi et al., 2017; Grigg and Hoffman, 1984; Lesniak et al., 2014). Values for SED were between 0.87 and 7.69 kPa, depending on the value for the inter-punctum interval used in the model (Table 1). Our estimates of SED were two to three orders of magnitude larger than that those reported by Elmi *et al*. (2017). There are two main factors contributing to this difference. First, SED is proportional to the Young’s modulus, which is 40 times larger in our simulations. Although the origin of this discrepancy is not clear, it may arise from differences in the finite element models used to infer the Young’s modulus. Second, different assumptions were used to deduce the minimum SED for single channel activation. Rather than assuming that maximal currents are produced by full gating of all the available channels (Elmi et al., 2017), we accounted for activation of both nearby and distant channels, many of which remained in a sub-conducting state (Fig. S1).

Because the SED estimate applies only to the responses to displacement steps and involves several uncertainties, we turned to our model linking body indentation to activation of single channels via elongation of a hypothetical gating filament. In our model (Sanzeni et al., 2018), the filament extension need to activate a single channel is of the order *g*_0_/ *g*_1_; such an elongation is generated by a deformation of constant rate *g*_0_/*g*_1_/*τ* if it is applied for a time of order, *τ*. Using values reported in Table 1, the model predicts that a single channel is able to sense a deformation on the order of 0.003 times the length of the linker generated over a time of 3 ms.

## Discussion

Using experiments and simulations, we advance understanding of how touch is transformed into depolarizing currents in touch receptor neurons. The picture emerging from this work is that touch sensitivity and filtering depend on a combination of factors: tissue mechanics and the distribution of ion channels within sensory neurons. These factors come together to generate mechanical forces transferred to individual channels whose activity accounts for the spatiotemporal dynamics of mechanoreceptor currents.

Additionally, our results show that the depth and speed of indentation affect the amplitude and time course of MRCs. Current-displacement curves do not saturate and instead display inflections that vary with internal pressure. By contrast, when we hold displacement constant, both the peak MRC-speed and charge-speed curves saturate at high speeds. The increase in peak current with speed occurs partially due to increased synchrony of channel openings with faster stimuli. Yet the total charge—which reflects all channel openings during the MRC—also increases with stimulus speed, suggesting that even without the effect of synchrony, MRCs depend on stimulus speed. This idea is also supported by the fact that ‘off’ responses, which have more synchronous activation, show a similar dependence on stimulus speed. Collectively, these observations indicate that activating MRCs in their native environment depends on tissue mechanics and dynamics as well as on the intrinsic mechanosensitivity of the MeT channels that carry the currents.

### Two mechanisms govern MRC amplitude by defining the strain field generated during touch

Displacement (indentation) and speed affect MRCs via two mechanisms: 1) recruiting individual channels; and 2) modulating the open probability for each channel. In general, the macroscopic current (*I*) is given by the product of the current carried by each individual channel (*i*), the number of channels available (*N*) and their probability of being open (*P*_*o*_). In many systems, we justifiably assume that *N* is independent of stimulus intensity. We can also assume that while *P*_*o*_ varies with stimulus intensity, it remains consistent across channels. For touch sensation, neither of these assumptions hold true. Our experimental and simulated data are consistent with the idea that larger displacements recruit increasingly distant channels by expanding the strain field. We previously showed that a displacement of a given magnitude elicits a larger response when the displacement begins from a pre-indented position (Sanzeni et al., 2018); this may occur due to a combination of wider recruitment and increased synchrony of activation. When we positioned the stimulus posterior to all channels, the current-displacement curve was shifted to the right (larger displacements) because we began to observe responses only once the displacement was large enough for the mechanical strain to reach the nearest channels (Figs. 4, 5, and 9). Our model also shows that both larger displacements and faster speeds increase the number of channels that fall within the indentation-induced strain field (Figs. 3D and 7F). Increasing either displacement or speed increases the probability that a channel will be in a sub-conducting or open state in a manner that depends on the channel’s distance from the stimulus (Fig. 9). The macroscopic current must take into account the different stimulus experienced by each channel as a result of tissue mechanics.

Further evidence that the joint effects of distal channel recruitment and modulation of open probability govern MRC amplitude comes from the finding that neither experimental nor simulated current-speed curves were well fit by the saturating Boltzmann function expected if open probability were the dominant factor affecting current size. The total charge, in fact, appeared to have two or more distinct steps that may reflect differences in the relative contributions of increased open probability and recruitment (Fig. 7B). Although we held the final displacement constant across speeds, the rate of strain, which determines the magnitude of the force acting on the channels in the model, increased with faster stimulation. Even if *N* remains constant across all speeds, the distribution of open probabilities likely changes with curvature and strain near the stimulator. Curvature and strain, in turn, depend on body mechanics. Our model predicts that channels distant from the stimulator position are more likely to be in a sub-conducting rather than a fully open state. Moreover, increasing stimulator speed might increase the proportion of fully open *vs.* sub-conducting channels in a non-linear manner. Collectively, these two mechanisms work together to expand the TRN operating range, enabling this neuron to detect stimuli which are small and fast as well as those which are slow and large.

How might nematodes leverage these two mechanisms of TRN activation? One possibility is to escape from predatory fungi that trap nematodes by means of constricting traps. These traps inflate within ∼100ms of contact (Higgins and Pramer, 1967), constricting to produce an initial indentation of roughly 15% of the worm’s body width (Barron, 1979; Nordbring-Hertz et al., 2011). This initial rapid, but modest stimulation may be followed by a slower but more complete closure of the constricting ring (Barron, 1979). Thus, the initial indentation would be expected to activate the touch receptor neurons swiftly while a larger indentation could recruit distal channels to ensure a robust escape if the initial response failed. Consistent with this idea, *C. elegans* that lack functional TRNs are more likely to be trapped by such fungi in the laboratory (Maguire et al., 2011).

### MRC kinetics and on/off asymmetry depend on channel recruitment

The model supports the idea that channels located farther away from the center of stimulation are recruited and opened later. However, while the rate at which the strain changes is as important for channel activation as the magnitude of the strain, the difference in timing is noticeable only at slower speeds (Fig. 9B). As seen in hair cell stereocilia (Ó Maoiléidigh and Ricci, 2019), distant channels experience increases in strain with a delayed time course because displacement along the worm is not instantaneous. The rate at which strain changes for distal channels can be high only when the displacement is large enough to reach them, which occurs late in an ‘on’ ramp or early in an ‘off’ ramp. Proximal channels, conversely, experience the greatest changes in strain at small displacements, where the curvature changes most drastically. As the proportion of distal to proximal channels increases with greater recruitment, the overall time course shifts toward the distal temporal profile, with slow ‘on’ activation and fast ‘off’ activation, creating asymmetry in MRC kinetics.

Similar to intact mammalian Pacinian corpuscles (Mendelson and Loewenstein, 1964), the DEG/ENaC/ASIC channels in *C. elegans* TRNs respond with depolarizing currents at both the onset and offset of an indentation step (Árnadóttir et al., 2011; Eastwood et al., 2015; O’Hagan et al., 2005). In this study, we also observed nearly identical responses at stimulus speeds around 1mm/s, but varying the speed varied the asymmetry between peak ‘on’ and ‘off’ currents. The significant increase in activation time and the relative invariance of total charge across speeds suggest that the high ‘off’/’on’ ratio at slow speeds is due to a change in synchrony (Fig. 7). Many MeT channels show asymmetrical responses to the onset and offset of a stimulus, due to inactivation (Lewis et al., 2017) or because the channels are part of asymmetrical structures that provide directional mechanical stimuli (Katta et al., 2015). Our current and previous models assume that individual channels receive symmetrical forces at the onset and offset of stimuli (Eastwood et al., 2015; Sanzeni et al., 2018). Here, we show that asymmetry of the stimulus is dynamic and is produced at the population level by the spatiotemporal recruitment of channels at different locations.

### Implications for understanding touch in vivo

The sensory neurons innervating Pacinian corpuscles show rapidly adapting, phasic responses similar to those of *C. elegans* mechanoreceptor neurons (Geffeney and Goodman, 2012). Adaptation of Pacinian corpuscle afferents slow following delamination of the corpuscle, indicating that the high-pass filter depends on the mechanics of surrounding tissues (Loewenstein and Mendelson, 1965; Loewenstein and Skalak, 1966). In addition, Pacinian corpuscles seem to require the lamellae to sustain the ‘off’ response. Although *C. elegans* TRNs differ in their anatomical structure, the mechanics of the body as a fluid-filled shell creates a similar mechanical filter that leads to adaptation. Furthermore, the congruence between model dynamics and experimental dynamics lends credence to the model. We can pursue this line of research by directly measuring the time course and distribution of strain throughout the body and along the TRN neurite. How would these MeT channels respond if we stimulated them in cultured neurons without body mechanics filtering the force? Would the channels still show the same rapidly-adapting behavior and responses to both stimulus onset and offset? The results presented here and in (Sanzeni et al., 2018) indicate that tissue mechanics play a crucial role in shaping the response to touch. Thus, we would not expect to recapitulate the dependence on indentation depth and speed *in vitro* that is found *in vivo*.

Although the model reflects and explains many of the dynamics we see experimentally without being tuned for this dataset, there are still differences that hint at biological mechanisms as yet unaccounted for by the model. While many of the quantitative discrepancies can be resolved by better matching existing parameters to the recordings, a few qualitative observations may require more consideration. For large displacements, the total charge for ‘on’ responses is consistently higher than the ‘off’ response at all speeds, which might suggest asymmetry in how force is transferred from the bead to the tissue during indentation vs. stimulus removal, inactivation at the level of the channel, or an irreversible dismantling of the molecules involved in transmitting force with repeated indentation. The presence of responses to distal posterior stimulation might be accounted for with simulation parameters tuned to this dataset, but it might also reflect our new observations about channel distribution. Perhaps the greater concentration of channels near the cell body allows the neuron to respond even when the strain field falls only partially on the neuron. Alternatively, extracellular collagen networks or cytoskeletal structures like microtubules could render the neuron stiffer than the surrounding tissue and direct force down the neurite. Luckily, *C. elegans* also provides many genetic tools for perturbing these elements and assessing their contributions to the spatial and temporal dynamics of mechanoreceptor responses. We can now explore the effects of mutations in skin and neuronal proteins on mechano-electrical transduction within the context of a larger, overarching biophysical model.

## Acknowledgements

We thank Z. Liao for superb technical support and animal husbandry; V. Vásquez, A. Peng, F. Loizeau and A. R. Ricci for assistance with the design, fabrication and testing of the mechanical stimulator and optical displacement sensor; and D. Ó Maoiléidigh and members of the Goodman laboratory for comments on the manuscript. Research supported by a Ruth L. Kirschstein fellowship (F31-NS-093825) to SK and NIH grants (R01-NS-047715; R35-NS-105092) to MBG.

## Author Contributions

All authors contributed to conceptualization, methodology, and revision of the final manuscript. Collection and analysis of electrophysiological data and indentation imaging data were performed by SK, while AD was responsible for analyzing the position of MEC-4 puncta, both with supervision by MBG. Simulations were carried out by AS, with supervision by MV. SK created the initial draft and visualizations.

## Abbreviations

SED: strain energy density
MeT: mechano-electrical transduction
MRC: mechanoreceptor current
TRN: touch receptor neuron

## Supplemental Figure Legend

**Figure S1.**
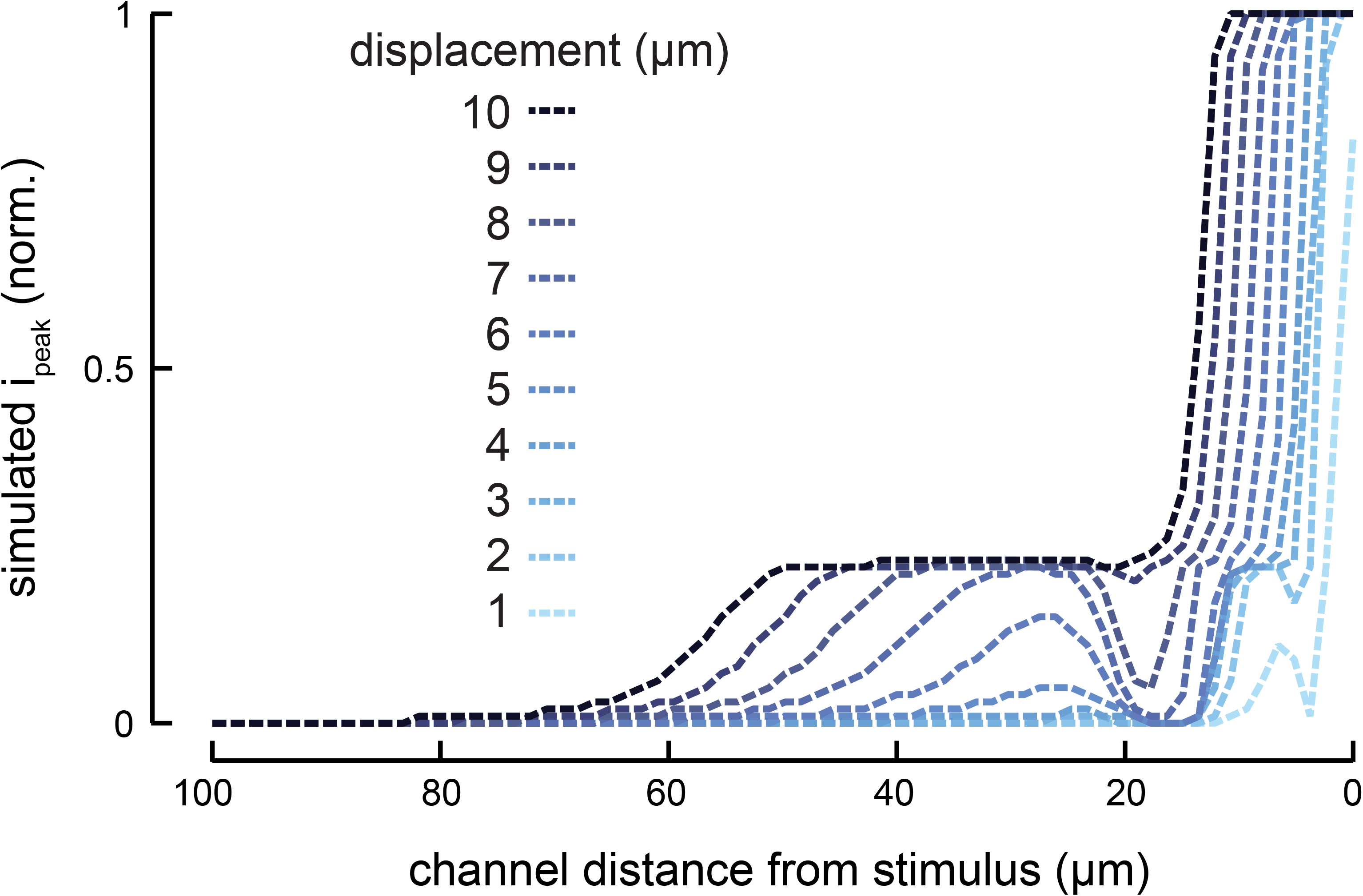
Individual channel responses depend on both channel distance and stimulus size. The peak current through individual channels (normalized to the single-channel open current *i*_*o*_ = 1.6 pA) in response to a step stimulus is highest for channels near the point of stimulation. At further distances, the current reflects the non-linearity of the strain field and increasing likelihood of the channel being in a sub-conducting rather than a fully open state.

## References

Árnadóttir, J., O’Hagan, R., Chen, Y., Goodman, M.B., Chalfie, M., 2011. The DEG/ENaC Protein MEC-10 Regulates the Transduction Channel Complex in Caenorhabditis elegans Touch Receptor Neurons. J. Neurosci. 31, 12695–12704. https://doi.org/10.1523/JNEUROSCI.4580-10.2011

Audoly, B., Pomeau, Y., 2010. Elasticity and Geometry: From hair curls to the nonlinear response of shells. Oxford University Press, Oxford, New York.

Barron, G.L., 1979. Observations on predatory fungi. Can. J. Bot. 57, 187–193. https://doi.org/10.1139/b79-028

Bekkers, J.M., Stevens, C.F., 1996. Cable properties of cultured hippocampal neurons determined from sucrose-evoked miniature EPSCs. J. Neurophysiol. 75, 1250–1255. https://doi.org/10.1152/jn.1996.75.3.1250

Bounoutas, A., O’Hagan, R., Chalfie, M., 2009. The Multipurpose 15-Protofilament Microtubules in C. elegans Have Specific Roles in Mechanosensation. Curr. Biol. 19, 1362–1367. https://doi.org/10.1016/j.cub.2009.06.036

Brown, A.L., Fernandez-Illescas, S.M., Liao, Z., Goodman, M.B., 2007. Gain-of-Function Mutations in the MEC-4 DEG/ENaC Sensory Mechanotransduction Channel Alter Gating and Drug Blockade. J. Gen. Physiol. 129, 161–173. https://doi.org/10.1085/jgp.200609672

Brown, A.L., Liao, Z., Goodman, M.B., 2008. MEC-2 and MEC-6 in the Caenorhabditis elegans Sensory Mechanotransduction Complex: Auxiliary Subunits that Enable Channel Activity. J. Gen. Physiol. 131, 605–616. https://doi.org/10.1085/jgp.200709910

Chalfie, M., Sulston, J., 1981. Developmental genetics of the mechanosensory neurons of Caenorhabditis elegans. Dev. Biol. 82, 358–370. https://doi.org/10.1016/0012-1606(81)90459-0

Chalfie, M., Thomson, J.N., 1979. Organization of neuronal microtubules in the nematode Caenorhabditis elegans. J. Cell Biol. 82, 278–289. https://doi.org/10.1083/jcb.82.1.278

Chelur, D.S., Ernstrom, G.G., Goodman, M.B., Yao, C.A., Chen, L., O’ Hagan, R., Chalfie, M., 2002. The mechanosensory protein MEC-6 is a subunit of the C. elegans touch-cell degenerin channel. Nature 420, 669–673. https://doi.org/10.1038/nature01205

Chen, X., Chalfie, M., 2015. Regulation of Mechanosensation in C. elegans through Ubiquitination of the MEC-4 Mechanotransduction Channel. J. Neurosci. 35, 2200–2212. https://doi.org/10.1523/JNEUROSCI.4082-14.2015

Cueva, J.G., Mulholland, A., Goodman, M.B., 2007. Nanoscale Organization of the MEC-4 DEG/ENaC Sensory Mechanotransduction Channel in Caenorhabditis elegans Touch Receptor Neurons. J. Neurosci. 27, 14089–14098. https://doi.org/10.1523/JNEUROSCI.4179-07.2007

Dandekar, K., Raju, B.I., Srinivasan, M.A., 2003. 3-D Finite-Element Models of Human and Monkey Fingertips to Investigate the Mechanics of Tactile Sense. J. Biomech. Eng. 125, 682–691. https://doi.org/10.1115/1.1613673

Eastwood, A.L., Sanzeni, A., Petzold, B.C., Park, S.-J., Vergassola, M., Pruitt, B.L., Goodman, M.B., 2015. Tissue mechanics govern the rapidly adapting and symmetrical response to touch. Proc. Natl. Acad. Sci. 112, E6955–E6963. https://doi.org/10.1073/pnas.1514138112

Edelstein, A., Amodaj, N., Hoover, K., Vale, R., Stuurman, N., 2010. Computer Control of Microscopes Using µManager. Curr. Protoc. Mol. Biol. 92, 14.20.1-14.20.17. https://doi.org/10.1002/0471142727.mb1420s92

Elmi, M., Pawar, V.M., Shaw, M., Wong, D., Zhan, H., Srinivasan, M.A., 2017. Determining the biomechanics of touch sensation in C. elegans. Sci. Rep. 7, 12329. https://doi.org/10.1038/s41598-017-12190-0

Emtage, L., Gu, G., Hartwieg, E., Chalfie, M., 2004. Extracellular Proteins Organize the Mechanosensory Channel Complex in C. elegans Touch Receptor Neurons. Neuron 44, 795–807. https://doi.org/10.1016/j.neuron.2004.11.010

Fehlauer, H., Nekimken, A.L., Kim, A.A., Pruitt, B.L., Goodman, M.B., Krieg, M., 2018. Using a Microfluidics Device for Mechanical Stimulation and High Resolution Imaging of C. elegans. JoVE J. Vis. Exp. e56530–e56530. https://doi.org/10.3791/56530

Geffeney, S.L., Cueva, J.G., Glauser, D.A., Doll, J.C., Lee, T.H.-C., Montoya, M., Karania, S., Garakani, A.M., Pruitt, B.L., Goodman, M.B., 2011. DEG/ENaC but Not TRP Channels Are the Major Mechanoelectrical Transduction Channels in a C. elegans Nociceptor. Neuron 71, 845–857. https://doi.org/10.1016/j.neuron.2011.06.038

Geffeney, S.L., Goodman, M.B., 2012. How We Feel: Ion Channel Partnerships that Detect Mechanical Inputs and Give Rise to Touch and Pain Perception. Neuron 74, 609–619. https://doi.org/10.1016/j.neuron.2012.04.023

Goodman, M.B., Hall, D.H., Avery, L., Lockery, S.R., 1998. Active Currents Regulate Sensitivity and Dynamic Range in C. elegans Neurons. Neuron 20, 763–772. https://doi.org/10.1016/S0896-6273(00)81014-4

Grigg, P., Hoffman, A.H., 1984. Ruffini Mechanoreceptors in Isolated Joint Capsule: Responses Correlated with Strain Energy Density. Somatosens. Res. 2, 149–162. https://doi.org/10.1080/07367244.1984.11800555

Higgins, M.L., Pramer, D., 1967. Fungal Morphogenesis: Ring Formation and Closure by Arthrobotrys dactyloides. Science 155, 345–346. https://doi.org/10.1126/science.155.3760.345

Hostettler, L., Grundy, L., Käser-Pébernard, S., Wicky, C., Schafer, W.R., Glauser, D.A., 2017. The Bright Fluorescent Protein mNeonGreen Facilitates Protein Expression Analysis In Vivo. G3 Genes Genomes Genet. 7, 607–615. https://doi.org/10.1534/g3.116.038133

Kang, L., Gao, J., Schafer, W.R., Xie, Z., Xu, X.Z.S., 2010. C. elegans TRP Family Protein TRP-4 Is a Pore-Forming Subunit of a Native Mechanotransduction Channel. Neuron 67, 381–391. https://doi.org/10.1016/j.neuron.2010.06.032

Katta, S., Krieg, M., Goodman, M.B., 2015. Feeling Force: Physical and Physiological Principles Enabling Sensory Mechanotransduction. Annu. Rev. Cell Dev. Biol. 31, 347–371. https://doi.org/10.1146/annurev-cellbio-100913-013426

Krieg, M., Stühmer, J., Cueva, J.G., Fetter, R., Spilker, K., Cremers, D., Shen, K., Dunn, A.R., Goodman, M.B., 2017. Genetic defects in β-spectrin and tau sensitize C. elegans axons to movement-induced damage via torque-tension coupling. eLife 6. https://doi.org/10.7554/eLife.20172

Lesniak, D.R., Gerling, G.J., 2009. Predicting SA-I mechanoreceptor spike times with a skin-neuron model. Math. Biosci. 220, 15–23. https://doi.org/10.1016/j.mbs.2009.03.007

Lesniak, D.R., Marshall, K.L., Wellnitz, S.A., Jenkins, B.A., Baba, Y., Rasband, M.N., Gerling, G.J., Lumpkin, E.A., 2014. Computation identifies structural features that govern neuronal firing properties in slowly adapting touch receptors. eLife 3, e01488. https://doi.org/10.7554/eLife.01488

Lewis, A.H., Cui, A.F., McDonald, M.F., Grandl, J., 2017. Transduction of Repetitive Mechanical Stimuli by Piezo1 and Piezo2 Ion Channels. Cell Rep. 19, 2572–2585. https://doi.org/10.1016/j.celrep.2017.05.079

Loewenstein, W.R., Mendelson, M., 1965. Components of receptor adaptation in a Pacinian corpuscle. J. Physiol. 177, 377–397. https://doi.org/10.1113/jphysiol.1965.sp007598

Loewenstein, W.R., Skalak, R., 1966. Mechanical transmission in a Pacinian corpuscle. An analysis and a theory. J. Physiol. 182, 346–378. https://doi.org/10.1113/jphysiol.1966.sp007827

Maguire, S.M., Clark, C.M., Nunnari, J., Pirri, J.K., Alkema, M.J., 2011. The C. elegans Touch Response Facilitates Escape from Predacious Fungi. Curr. Biol. 21, 1326–1330. https://doi.org/10.1016/j.cub.2011.06.063

Mendelson, M., Loewenstein, W.R., 1964. Mechanisms of Receptor Adaptation. Science 144, 554–555. https://doi.org/10.1126/science.144.3618.554

Nekimken, A.L., Fehlauer, H., Kim, A.A., Manosalvas-Kjono, S.N., Ladpli, P., Memon, F., Gopisetty, D., Sanchez, V., Goodman, M.B., Pruitt, B.L., Krieg, M., 2017a. Pneumatic stimulation of C. elegans mechanoreceptor neurons in a microfluidic trap. Lab. Chip 17, 1116–1127. https://doi.org/10.1039/C6LC01165A

Nekimken, A.L., Mazzochette, E.A., Goodman, M.B., Pruitt, B.L., 2017b. Forces applied during classical touch assays for Caenorhabditis elegans. PLOS ONE 12, e0178080. https://doi.org/10.1371/journal.pone.0178080

Nordbring-Hertz, B., Jansson, H.-B., Tunlid, A., 2011. Nematophagous Fungi, in: ELS. American Cancer Society. https://doi.org/10.1002/9780470015902.a0000374.pub3

Ó Maoiléidigh, D., Ricci, A.J., 2019. A Bundle of Mechanisms: Inner-Ear Hair-Cell Mechanotransduction. Trends Neurosci. 42, 221–236. https://doi.org/10.1016/j.tins.2018.12.006

O’Hagan, R., Chalfie, M., Goodman, M.B., 2005. The MEC-4 DEG/ENaC channel of Caenorhabditis elegans touch receptor neurons transduces mechanical signals. Nat. Neurosci. 8, 43–50. https://doi.org/10.1038/nn1362

Peng, A.W., Ricci, A.J., 2016. Glass Probe Stimulation of Hair Cell Stereocilia, in: Sokolowski, B. (Ed.), Auditory and Vestibular Research: Methods and Protocols. Springer New York, New York, NY, pp. 487–500. https://doi.org/10.1007/978-1-4939-3615-1_27

Petzold, B.C., Park, S.-J., Mazzochette, E.A., Goodman, M.B., Pruitt, B.L., 2013. MEMS-based force-clamp analysis of the role of body stiffness in C. elegans touch sensation. Integr. Biol. 5, 853–864. https://doi.org/10.1039/C3IB20293C

Quindlen, J.C., Lai, V.K., Barocas, V.H., 2015. Multiscale Mechanical Model of the Pacinian Corpuscle Shows Depth and Anisotropy Contribute to the Receptor’s Characteristic Response to Indentation. PLOS Comput. Biol. 11, e1004370. https://doi.org/10.1371/journal.pcbi.1004370

Quindlen-Hotek, J.C., Barocas, V.H., 2018. A finite-element model of mechanosensation by a Pacinian corpuscle cluster in human skin. Biomech. Model. Mechanobiol. 17, 1053–1067. https://doi.org/10.1007/s10237-018-1011-1

Rall, W., 1967. Distinguishing theoretical synaptic potentials computed for different soma-dendritic distributions of synaptic input. J. Neurophysiol. 30, 1138–1168. https://doi.org/10.1152/jn.1967.30.5.1138

Rivera Gomez, K.A., Schvarzstein, M., 2018. Immobilization of nematodes for live imaging using an agarose pad produced with a Vinyl Record. MicroPublication Biol.

Sanzeni, A., Katta, S., Petzold, B., Pruitt, B.L., Goodman, M.B., Vergassola, M., 2018. Tissue mechanics and somatosensory neural responses govern touch sensation in C. elegans. bioRxiv 471904. https://doi.org/10.1101/471904

Suzuki, H., Kerr, R., Bianchi, L., Frøkjær-Jensen, C., Slone, D., Xue, J., Gerstbrein, B., Driscoll, M., Schafer, W.R., 2003. In Vivo Imaging of C. elegans Mechanosensory Neurons Demonstrates a Specific Role for the MEC-4 Channel in the Process of Gentle Touch Sensation. Neuron 39, 1005–1017. https://doi.org/10.1016/j.neuron.2003.08.015

Thein, M.C., McCormack, G., Winter, A.D., Johnstone, I.L., Shoemaker, C.B., Page, A.P., 2003. Caenorhabditis elegans exoskeleton collagen COL-19: An adult-specific marker for collagen modification and assembly, and the analysis of organismal morphology. Dev. Dyn. 226, 523–539. https://doi.org/10.1002/dvdy.10259

Vásquez, V., Krieg, M., Lockhead, D., Goodman, M.B., 2014. Phospholipids that Contain Polyunsaturated Fatty Acids Enhance Neuronal Cell Mechanics and Touch Sensation. Cell Rep. 6, 70–80. https://doi.org/10.1016/j.celrep.2013.12.012

